# Inhibition of Metastatic Uveal Melanoma Cell Proliferation by the Histone Deacetylase 6 Inhibitor, Ricolinostat, is Linked to Altered Microphthalmia-associated Transcription Factor and Phospho-ERK Expression

**DOI:** 10.1101/2021.10.28.466226

**Authors:** Husvinee Sundaramurthi, Sandra García-Mulero, Kayleigh Slater, Simone Marcone, Josep M. Piulats, R. William Watson, Desmond J. Tobin, Lasse D. Jensen, Breandán N. Kennedy

## Abstract

Metastatic uveal melanoma (MUM) is characterized by poor patient survival. Unfortunately, current treatment options demonstrate limited benefits. In this study, we evaluate the efficacy of ACY-1215, a histone deacetylase 6 inhibitor (HDAC6i), to attenuate MUM cell growth *in vitro* and *in vivo,* and elucidate the underlying molecular mechanisms. Treatment of OMM2.5 MUM cells with ACY-1215 resulted in a significant (*p* = 0.0001), dose-dependent reduction in cell survival and proliferation *in vitro*, and *in vivo* regression of primary OMM2.5 xenografts in zebrafish larvae. Furthermore, flow cytometry analysis revealed that ACY-1215 significantly arrested the OMM2.5 cell cycle in S phase (*p* = 0.0006) following 24 hours of treatment and significant apoptosis was triggered in a time- and dose-dependent manner (*p* = <0.0001). Additionally, ACY-1215 treatment resulted in a significant reduction in OMM2.5 p-ERK expression levels. Through proteome-profiling, attenuation of the microphthalmia-associated transcription factor (MITF) signaling pathway was linked to the observed anti-cancer effects of ACY-1215. In agreement, pharmacological inhibition of MITF signaling with ML329, significantly reduced OMM2.5 cell survival and viability *in vitro* (*p* = 0.0001) and *in vivo* (*p* = 0.0006). Our findings provide evidence that ACY-1215 and ML329 are efficacious against growth and survival of MUM cells and are potential therapeutic options for MUM.

**Simple Summary:** Uveal melanoma (UM) is the most common adult eye cancer. UM originates in the iris, ciliary body, or choroid (collectively known as the uvea), in the middle layer of the eye. This first or primary UM is treated by targeting the cancer cells using ocular radiation implants or by surgical removal of the eye. However, when UM spreads to the liver and other parts of the body, patients have a poor survival prognosis. Unfortunately, there are no effective treatment options for UM that has spread. Our aim is to help identify effective treatments for UM cancer that has spread. In our study, we identified that the drug ACY-1215 prevents the growth of UM cells from the liver. Our study has found a promising treatment approach for advanced UM.

## Introduction

Uveal melanoma (UM) is the most common adult intraocular cancer, afflicting approximately 4.3 per million people worldwide [1]. Although a rare cancer, incidence rates increase geographically in a South to North gradient, with countries like Ireland, Norway and Denmark reported to have the highest incidences in Europe [2–4]. UM originates with different frequencies in the uveal tract of the eye: choroid (∼90%), iris (∼4%) and ciliary body (∼6%) [5]. The most effective treatments for primary UM include surgical resection of the tumor, radiotherapy (plaque brachytherapy or proton-beam therapy) and enucleation of the affected eye [4, 6]. Unfortunately, approximately 50% of patients diagnosed with primary UM progress to develop metastatic UM (MUM), primarily in the liver (∼89%), which is associated with poor survival prognosis (median overall survival (OS) ranging from 4 to 15 months) [2,7,8]. There is no *standard of care* treatment for MUM patients and current therapeutic options have limited benefit. MUM patients receive site -directed therapies (including surgical resection of tumor), the chemotherapeutic drug Dacarbazine (commonly used to treat cutaneous melanoma) or immunotherapy drugs such as Ipilimumab and Pembrolizumab [8, 9]. Unfortunately, treatment with Dacarbazine either as a monotherapy or combinatorial therapy did not improve overall survival or progression free survival [8,10–12]. In addition, immune checkpoint inhibitors, MEK inhibitors and liver-directed therapies are in clinical and preclinical trials for UM at present [13–15]. Recently, a Phase III clinical study with Tebentafusp (a form of immunotherapy that recruits and redirects T cells to tumor cells) reported favorable evidence in MUM patients with the one-year OS rate reported at 73% (N = 252) in the Tebentafusp treatment group compared to 59% (N = 126) in the control group, with an estimated median OS of 21.7 months and 16.0 months, respectively [16]. Nevertheless, there is still an imperative to identify highly efficacious novel drugs for the treatment of MUM, as Tebentafusp has only been trialed in a subset of MUM patient cohort who are HLA-A*02:01-positive.

Histone deacetylase inhibitors (HDACi) have garnered widespread interest in the past two decades as anti- cancer agents [17–21]. Four pan-HDAC inhibitors - Vorinostat (SAHA) for relapsed and refractory cutaneous T-cell lymphoma (CTCL), Belinostat for peripheral T-cell lymphoma (PTCL), Romidepsin for CTCL/PTCL and Panobinostat for multiple myeloma, are approved for market use as treatment options by the FDA and/or EMA [18]. Chidamide is approved by the Chinese FDA for treatment of PTCL, with more research underway in other cancers [22]. Pre-clinical studies identified pan-HDACi to show efficacy as anti-cancer agents in UM and/or MUM cell lines, *in vitro* and/or *in vivo* [23]. Encouragingly, the first Phase II clinical trial with 29 MUM patients reported that a combination treatment of Entinostat (pan-HDACi) and Pembrolizumab (PD-1 inhibitor) resulted in a median OS of 13.4 months with one year OS reported as 59%; and median progression free survival (PFS) of 2.1 months and a 17% one year PFS [24, 25]. More recently, a novel compound, VS13, which displays increased selectivity against HDAC6, reduced UM cell viability [26].

In relation to selective HDAC inhibition, histone deacetylase 6 inhibitors (HDAC6i) show promise as anti- cancer agents in preclinical studies; and are currently under clinical trial investigations as a monotherapy or combinatorial therapy for lymphoproliferative disease, hematologic malignancies, and solid tumors [27–30]. HDAC6 is a Class IIb enzyme and unlike other HDAC isozymes, mainly resides in the cytoplasm and acts primarily on cytosolic proteins [31]. This provides a potential selective advantage over pan-HDAC inhibitors due to their pleiotropic effects. Pre-clinical studies report multiple selective HDAC6i compounds as anti-cancer agents with anti-cell proliferation, anti-cell viability, and tumor attenuation in glioblastoma, ovarian cancer, and bladder cancer [17,30,32–34]. A handful of HDAC6i clinical trials are registered and currently proceeding. A Phase Ib/II trial of ACY-1215 (Ricolinostat) in a small cohort of lymphoma patients revealed it was well-tolerated, and the disease stabilized in 50% (8 out of 16 patients evaluated) of patients [27]. ACY-1215 in combination with paclitaxel, was well tolerated and exhibited activity in patients with ovarian cancer in a small-scale Phase Ib trial, which was prematurely terminated [28]. In a Phase I/II trial in patients with relapsed or refractory multiple myeloma, ACY-1215 given in combination with Bortezomib and Dexamethasone, was well tolerated and active as an anti-myeloma agent [29]. There are also ongoing clinical trials with HDAC6 inhibitors (*e.g.,* ACY-1215, Citarinostat (ACY-241) or KA2507) as a single agent or combination therapy for non-small cell lung cancer, metastatic breast cancer and solid tumor [30].

Here, we investigated the efficacy of HDAC6i, specifically ACY-1215, to inhibit MUM cell growth *in vitro* and *in vivo*; and to understand the molecular mechanism of ACY-1215 in MUM cell survival and proliferation. ACY-1215 significantly attenuated growth of MUM cells *in vitro* and *in vivo*; and this effect is correlated to regulation of microphthalmia-associated transcription factor (MITF) and phospho-ERK (p- ERK) expression levels, which offers additional therapeutic targets for MUM.

## Results

### ACY-1215 significantly attenuates long term proliferation of human uveal melanoma cell lines

Three commercially available HDAC6i (Tubastatin A, ACY-1215 and Tubacin) were selected to determine their efficacy in reducing long-term proliferation of human UM cell lines derived from primary (Mel285 and Mel270) and metastatic (OMM2.5) UM tumors [35-37]. Cells were treated for 96 hours at selected concentrations, treatment was stopped, cells were cultured for another 10 days in fresh complete media and colonies formed visualized with crystal violet staining and counted [38]. Initial screens at 10 - 50 µM showed a dose-dependent reduction in UM cell proliferation with all three HDAC6i tested **(Figure S1)**. ACY-1215 was selected as the highest-ranked drug for subsequent studies based on its observed effects in all three UM cell lines tested and its existing approved use in clinical trials [27–29]. ACY-1215 was tested at 1, 5, 10, 20 and 50 µM concentrations in Mel270 and OMM2.5 cells, as these cell lines were established from the same patient **(Figure 1)**. Mel270 cells showed significant reductions in viable clones averaging 94.7%, 99.98%, 99.98% and 99.8% (*p* = 0.0001) decreases at 5, 10, 20 and 50 µM concentration of ACY- 1215, respectively, compared to vehicle controls (0.5% DMSO) **(Figure 1B and 1C)**. Similarly, in OMM2.5 cells, ACY-1215 significantly reduced surviving colonies in a dose-dependent manner averaging 92.9%, 99.5%, 99.98% and 99.8% (*p* = 0.0001) decreases when treated at 5, 10, 20 and 50 µM respectively, compared to vehicle control **(Figure 1B and 1D)**. Patients diagnosed with MUM are previously prescribed the chemotherapeutic Dacarbazine, hence this was used as a clinical control treatment on both Mel270 and OMM2.5 cells. However, there was no significant difference observed at the tested concentration of 20 µM in either Mel270 (9.8% increase in colony formation, *p* = 0.095) or OMM2.5 (16.5% increase in colony formation, *p* = 0.704) cells **(Figure 1B, 1C and 1D)**. As our primary goal was to identify novel treatment strategies for MUM, follow-on studies determined ACY-1215 efficacy *in vivo* and mechanism of action in OMM2.5 cells.

**Figure 1:**
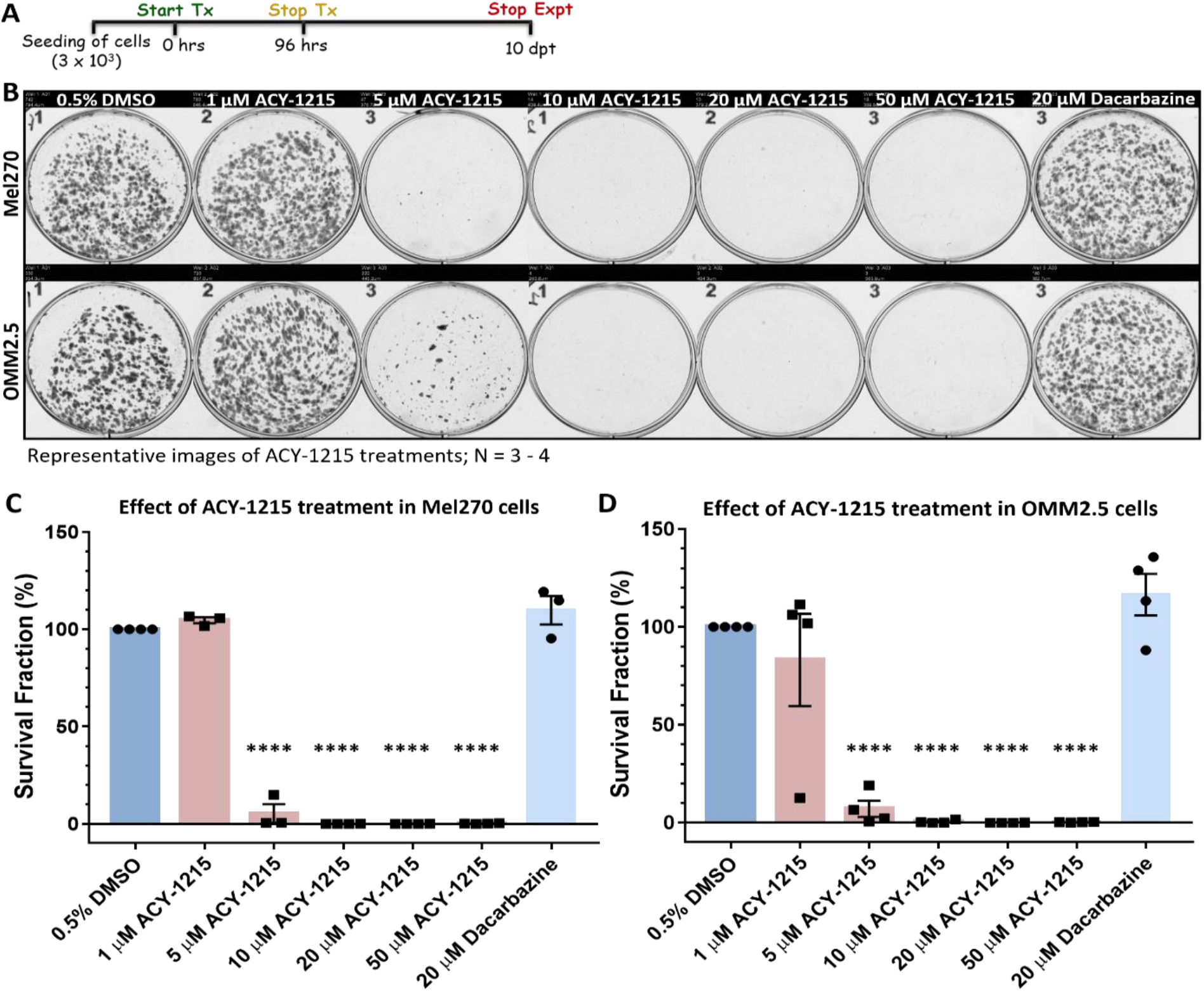
ACY-1215 is efficacious as an anti-cancer drug in UM and MUM cell lines. A: Schematic diagram on the treatment regime followed. B: Representative image of clonogenic assay plates for Mel270 (top panel) and OMM2.5 cells (bottom panel) treated with 0.5% DMSO; 1, 5, 10, 20 or 50 μM ACY-1215 or 20 μM Dacarbazine for 96 hours. C and D: A dose-dependent, significant decrease in the surviving number of OMM2.5 colonies was observed, indicative of reduction in cell viability upon ACY-1215 treatment in comparison to 0.5% DMSO treatment. One-way ANOVA with Dunnett’s Test for Multiple Comparisons statistical analysis was performed, error bars represent mean ± SEM, \*\*\*\**p* value of 0.0001 (N = 3 - 4).

### Zebrafish OMM2.5 xenografts proved that ACY-1215 is efficacious in vivo

Our *in vitro* study provided preliminary evidence that ACY-1215 has anti-UM properties. Therefore, the efficacy of ACY-1215 *in vivo* was evaluated using a pre-clinical model of MUM, zebrafish OMM2.5 xenografts. A toxicity screen determined the maximum tolerated dose of ACY-1215 and Dacarbazine in zebrafish larvae, with both drugs well-tolerated at all tested concentrations **(Figure S2)**. OMM2.5 Dil labelled cells were transplanted into the perivitelline space of 2 days old larvae and xenografts were treated with 0.5% DMSO, 20 µM ACY-1215 or 20 µM Dacarbazine for 3 days (5 days old) **(Figure 2A)**. These concentrations were selected based on the *in vitro* studies conducted. Primary xenograft fluorescence from OMM2.5 transplants regressed by approximately 65% (*p* = <0.0001) with 20 µM ACY-1215 treatment compared to vehicle controls **(Figure 2B and 2D)**. There was no notable difference in primary xenograft fluorescence when treated with 20 µM Dacarbazine in comparison to vehicle control. Additionally, the ability of transplanted OMM2.5 cells to disseminate was assessed by the number of cells present at the caudal vein plexus, 3 days post treatment. Dissemination of OMM2.5 Dil labelled cells was not affected by either 20 µM ACY-1215 or 20 µM Dacarbazine **(Figure 2C and 2E)**. On average, four disseminated OMM2.5 Dil labelled cells were detected at the caudal vein plexus of ACY-1215 treated larvae and five disseminated cells were counted in larvae treated with either 20 µM Dacarbazine or 0.5% DMSO. In summary, ACY-1215 at the tested concentration is effective in preventing UM cell growth/viability but not dissemination of OMM2.5 xenografts, *in vivo*.

**Figure 2:**
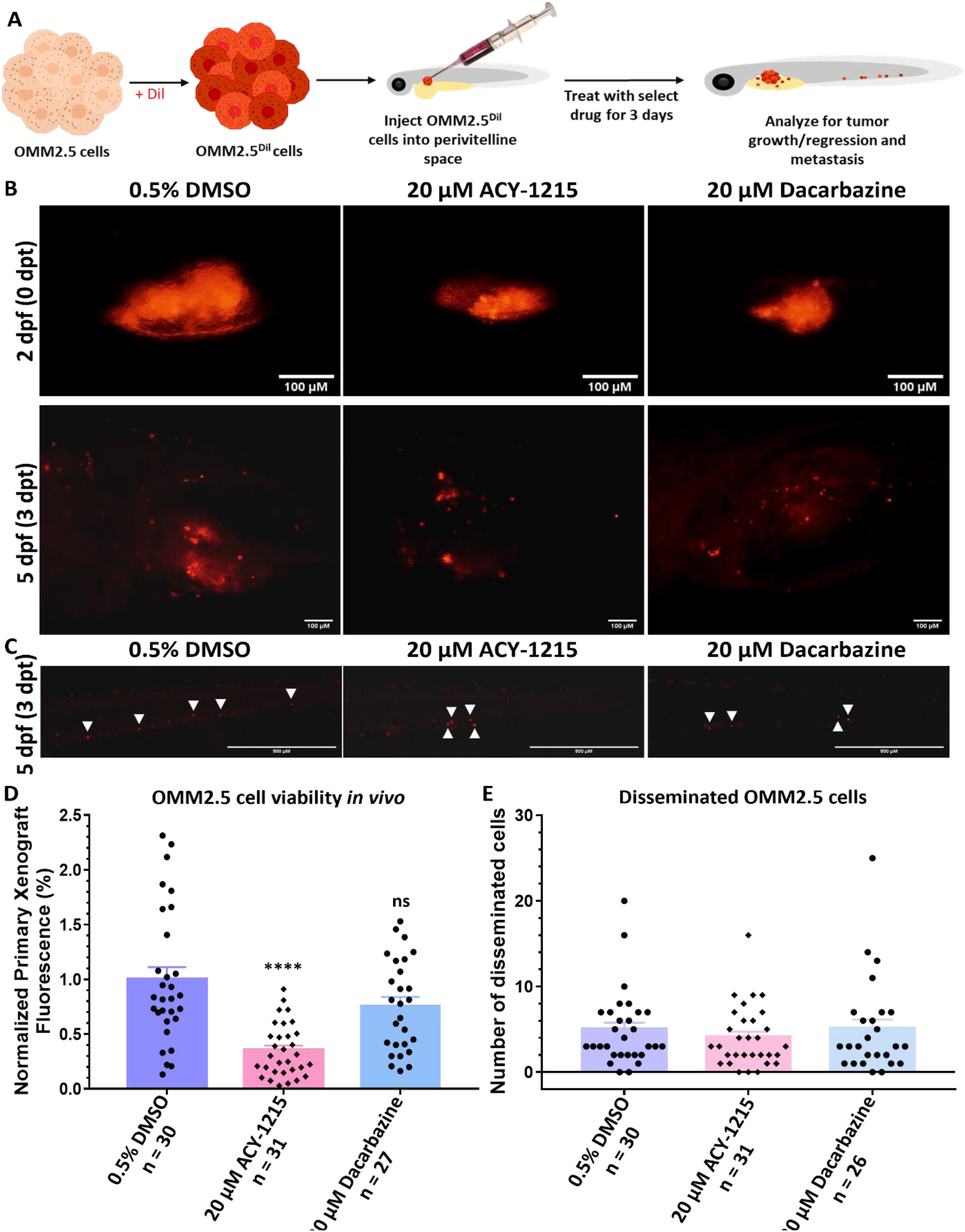
ACY-1215 demonstrates anti-cancer effects *in vivo* in zebrafish OMM2.5 xenografts. A: Schematic depicting the workflow for assessing ACY-1215 effects *in vivo*. B: Top panel shows representative images of transplanted OMM2.5 Dil labelled cells into the perivitelline space of 2 days old zebrafish larvae. Bottom panels present representative images of the distribution of OMM2.5 Dil labelled cells in xenografts 3 days post treatment (dpt) with 0.5% DMSO (n = 30), 20 μM ACY-1215 (n = 31) or 20 μM Dacarbazine (n = 27). C: At 3 dpt, OMM2.5 Dil labelled cells have disseminated (white arrowhead) to the caudal vein plexus of the zebrafish larvae. D: ACY-1215 treatment for 3 days, resulted in a significant (****, *p* = 0.0001) reduction in normalized primary xenograft fluorescence on average in comparison to larvae treated with 0.5% DMSO or 20 μM Dacarbazine. E: No difference observed in the average number of disseminated cells between vehicle control treated or drug-treated groups after 3 days. Statistical analysis was performed using One-way ANOVA with Dunnett’s Test for Multiple Comparisons and error bars present mean ± SEM.

### Analysis of ACY-1215 Targets in UM Patients Samples and UM cells

HDAC6 is a selective target of ACY-1215 at lower concentrations, hence HDAC6 expression in the different UM/MUM cell lines was confirmed by immunoblotting (**Figure 3**). No significant difference in HDAC6 expression was detected when the untreated primary ocular tumor derived cell lines (Mel270 and Mel285) or untreated MUM (OMM2.5) cell line were compared to untreated ARPE19 cells, a human retinal pigment epithelium cell line **(Figure 3A, 3A’ and Figure S3)**. To determine if ACY-1215 was indeed blocking HDAC6 activity, expression of its downstream substrate, acetylated α-tubulin was analyzed [30]. We observed a significant increase in acetylated α-tubulin levels after 4 (3.56-fold increase, *p* = 0.001) and 24 (3.67-fold increase, *p* = 0.0002) hours post treatment (hpt) with 20 µM ACY-1215 compared to 0.5% DMSO treated OMM2.5 cells, confirming the inhibitory effects of ACY-1215 **(Figure 3B, 3B’ and Figure S7A)**. As a dose-dependent anti-cancer effect of ACY-1215 was observed in the clonogenic assays and zebrafish xenografts, correlations between expression level of *HDAC6* and UM patient overall survival/progression free survival was analyzed. Extracting the gene expression data of 80 primary UM samples from The Cancer Genome Atlas (TCGA), Cox proportional-hazards models and Kaplan-Meier survival curves were generated. Kaplan-Meier survival curves were generated with a cut-off of 50% to demarcate as high or low *HDAC6* expression, and Log-rank test was used to compare survival probability between groups. Interestingly, high *HDAC6* expression was significantly associated with better overall survival but not with progression free survival (Cox OS, *p* = 0.007 and Cox PFS, *p* = 0.154) **(Figure 3C)**.

**Figure 3:**
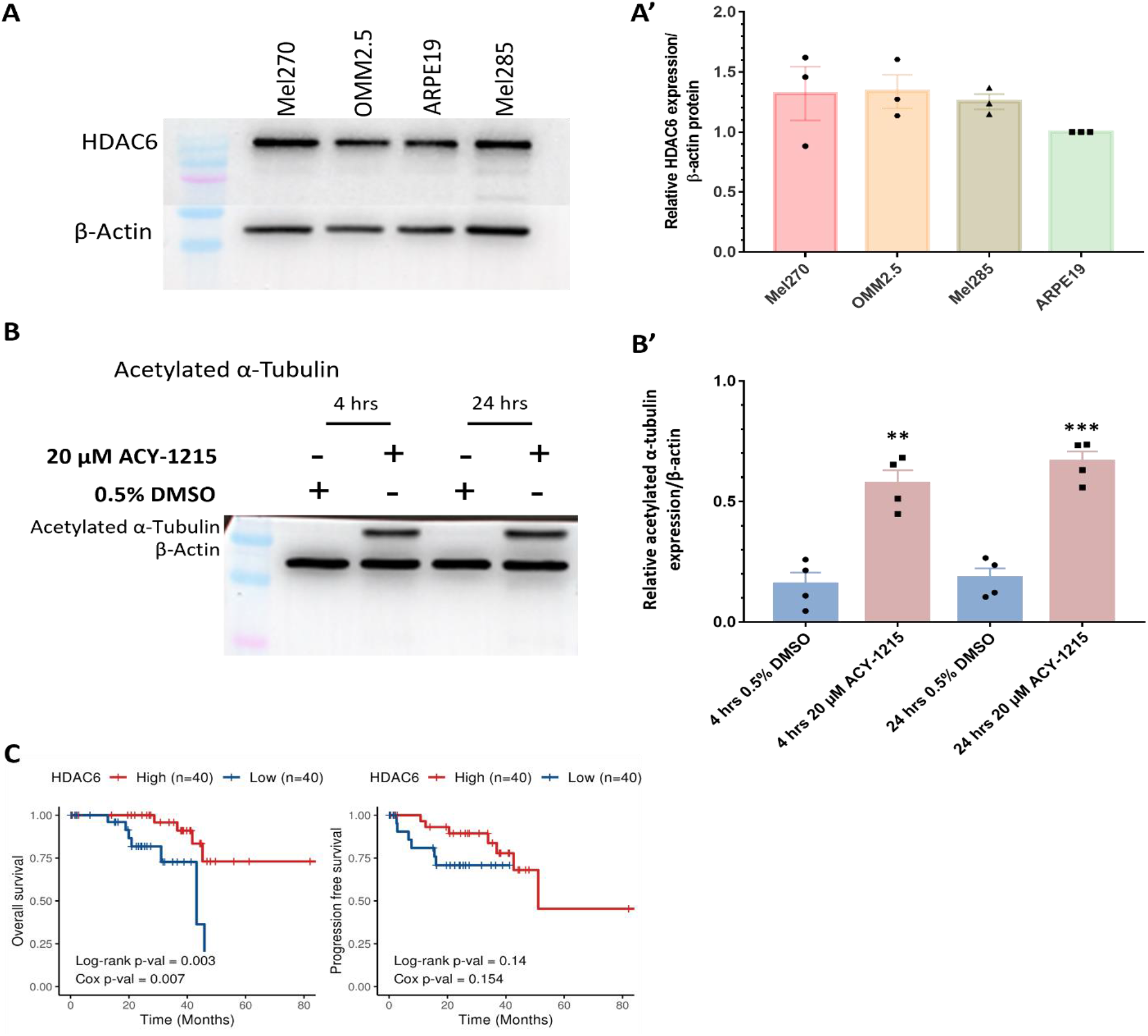
Expression and activity of HDAC6 in UM/MUM cells. A and A’: HDAC6 is expressed in Mel270, OMM2.5, Mel285 and ARPE19 cells (N = 3). B and B’: 20 μM ACY-1215 treatment significantly increased acetylated α-tubulin expression levels at 4 hpt (**, *p* = 0.0013) and 24 hpt (***, *p* = 0.0002) compared to 0.5% DMSO treatment. Student’s Unpaired T test statistical analysis was performed, and data presented as mean ± SEM. Representative blots for each protein probed and densitometry analysis presented, plus raw blots are provided in Supplementary Figures 3 and 7. C: Kaplan-Meier survival curves demonstrating correlation between expression of HDAC6 and overall survival (OS) or progression free survival (PFS) in UM patients. Median values were used as cut-off for high (red) and low (blue) expression levels, with Log-rank *p*-values (categorical variable) and Cox *p*-values (continuous variable) calculated (n = 80).

A known caveat of ACY-1215 is the non-selective inhibition of other HDAC isozymes at higher concentrations. The reported IC50 of ACY-1215 is 4.7 nM, at which ACY-1215 acts as a highly potent and selective HDAC6 inhibitor. Hence, we postulated that the observed effects of ACY-1215 in OMM2.5 cells are partly attributed to parallel inhibition of other HDACs. At higher concentrations, ACY-1215 inhibits HDAC 2, 3, 1, 8, 7, 5, 4, 9, 11 and SIRT 1/2 **(Figure S4A and S4B)** [39]. Thus, correlations between these HDAC isoforms and UM OS/PFS probability was analyzed **(Figure S4C)**. *HDAC2* (Cox OS, *p* = 0.1; Cox PFS, *p* = 0.454), *HDAC3* (Cox OS, *p* = 0.443; Cox PFS, *p* = 0.293) and *HDAC1* (Cox OS, *p* = 0.219; Cox PFS, *p* = 0.408) expression does not correlate with OS or PFS, respectively. Intriguingly, high *HDAC11* expression correlated significantly with better OS and PFS (Cox OS, *p* = 0.006; Cox PFS, *p* = 0.024). On the other hand, *HDAC8* (Cox OS, *p* = 0.231), *HDAC7* (Cox OS, *p* = 0.751), *HDAC5* (Cox OS, *p* = 0.837), *HDAC4* (Cox OS, *p* = 0.34), *HDAC9* (Cox OS, *p* = 0.704) and *SIRT1* (Cox OS, *p* = 0.579) expression did not significantly correlate to overall survival probability. Low expression of *HDAC8* (Cox PFS, *p* = 0.024), *HDAC7* (Cox PFS, *p* = 0.05), *HDAC5* (Cox PFS, *p* = 0.012), *HDAC4* (Cox PFS, *p* = 0.012), *HDAC9* (Cox PFS, *p* = 0.00001) and *SIRT1* (Cox PFS, *p* = 0.023) significantly correlated to a better PFS probability. There was significant correlation between high *SIRT2* expression and OS probability (Cox OS, *p* = 0.025) while its expression did not correlate to PFS (Cox PFS, *p* = 0.531). In summary, HDAC6 expression levels were not altered across the three UM/MUM cell lines analyzed, and high *HDAC6* expression level is associated with better survival for UM patients.

### Proteome profiling uncovers molecular signals altered in OMM2.5 UM cells by ACY-1215

Having observed beneficial effects against the growth and viability of UM cell lines *in vitro* and *in vivo*, proteome profiling of ACY-1215 treated OMM2.5 cells was performed to investigate the molecular mechanism of its anti-cancer action **(Figure 4, Figure S5, Table S1 and S2)**. Changes in protein expression levels were analyzed after 4 and 24 hours of 20 µM ACY-1215 treatment **(Figure 4A)**. A total of 4,423 proteins were detected across all samples by mass spectrometry. At 4 hpt, 42 proteins were differentially expressed with 11 proteins significantly upregulated and 30 proteins significantly downregulated **(Figure S5A and S5B)**. Using Cluego pathway analysis, the terms *dendrite development* and *regulation of G protein-coupled receptor signaling* pathways were identified as downregulated **(Figure S5C)**. A distinct pathway was not detected within the upregulated proteins. At 24 hpt, 150 proteins and 202 proteins were significantly down- and up-regulated, respectively **(Figure 4B)**. GO pathway enrichment analysis (fold change of ≥ 1.2) for biological processes identified multiple pathways downregulated by ACY-1215, with *pigment granule organization* (11.24% of proteins) and *pigment cell differentiation* (7.87% of proteins) being prominently altered **(Figure 4C, Figure S6A and Table S1)**. Through enriched pathway analysis, biological processes such as *regulation of microtubule polymerization* or *depolymerization* (7.25% of proteins), *DNA duplex unwinding* (3.11% of proteins), *regulation of chromatin silencing* (3.11% of proteins), *regulation of extrinsic apoptotic signaling pathway in absence of ligand* (2.07 % of proteins), *cellular senescence* (1.55% of proteins), *exit from mitosis* (1.55% of proteins) and *ERBB2 signaling pathway* (1.55% of proteins) were significantly upregulated by ACY-1215 treatment in OMM2.5 cells **(Figure 4D, Figure S6B and Table S2)**. Proteins Arginase-2, mitochondrial (ARG2; 13.05-fold), Semenogelin-2 (SEMG2; 10.26-fold), Protein AHNAK2 (AHNAK2; 8.69-fold), Neurosecretory protein VGF (VGF; 7.29-fold), Nuclear receptor subfamily 4 group A member 1 (NR4A1; 6.86-fold), Thymidine kinase, cytosolic (TK1; 5.72-fold), PRKC apoptosis WT1 regulator protein (PAWR; 4.80-fold), Tudor and KH domain containing, isoform CRA_a (TDRKH; 4.48-fold), Bromodomain-containing protein 2 (BRD2; 3.91-fold) and Ubiquitin-conjugating enzyme E2 S (UBE2S; 3.85-fold) were within the top ten significantly upregulated proteins **(Figure 5A)**.

**Figure 4:**
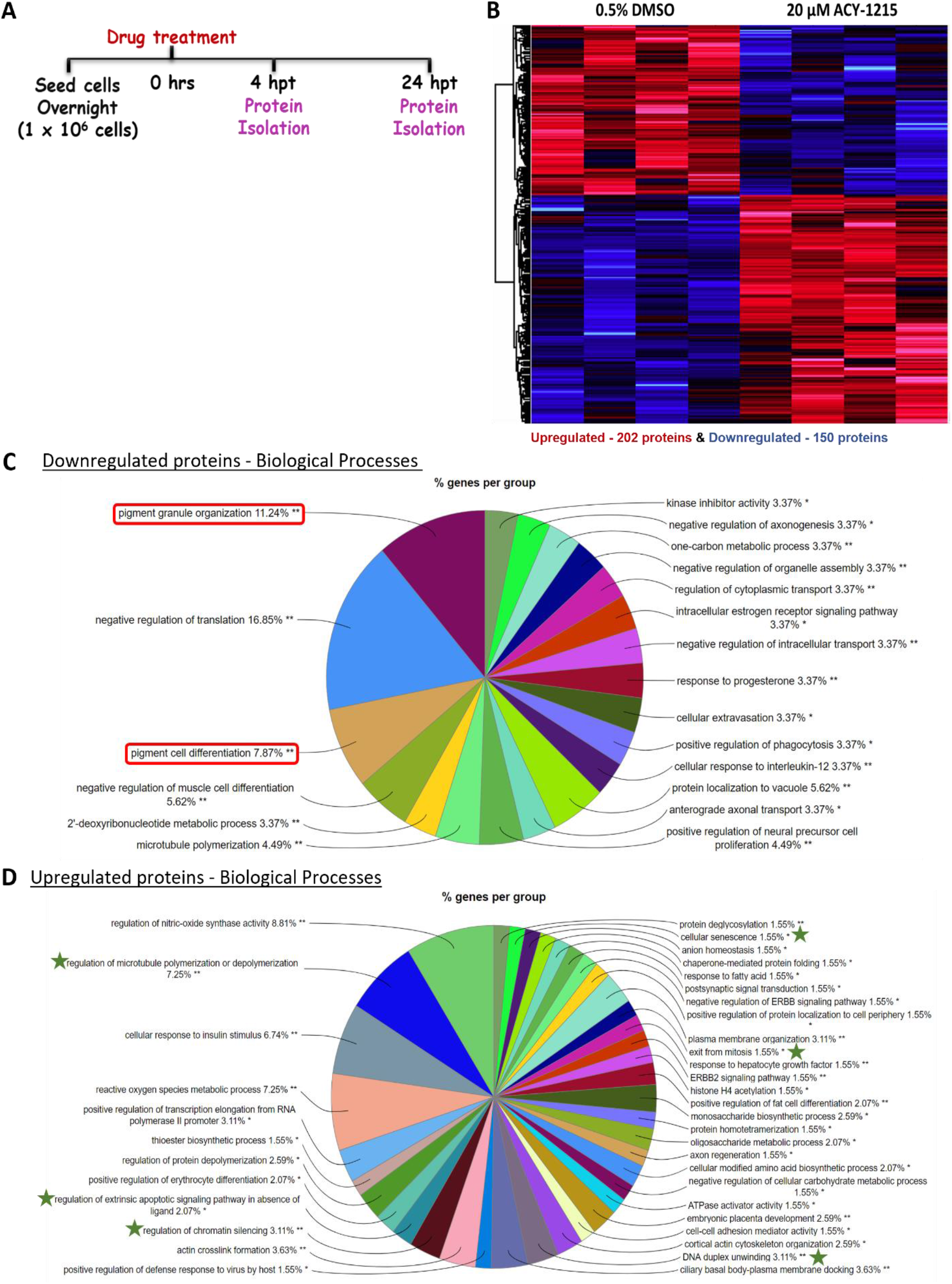
Proteome profiling of ACY-1215 treated cells to uncover mechanism of action. A: ACY-1215 treatment regime for proteome profiling of OMM2.5 cells. B: Heat map showing all significant differentially expressed proteins at 24 hours post 20 μM ACY-1215 treatment. A total of 4423 proteins were identified in MS with 150 downregulated (blue) and 202 upregulated (red) proteins (N = 4). C and D: Enriched protein pathway analysis for GO term: biological processes, for down- and up-regulated proteins given a fold change cut off of +/- ≥ 1.2, *p* ≤ 0.05 displayed as pie charts.

**Figure 5:**
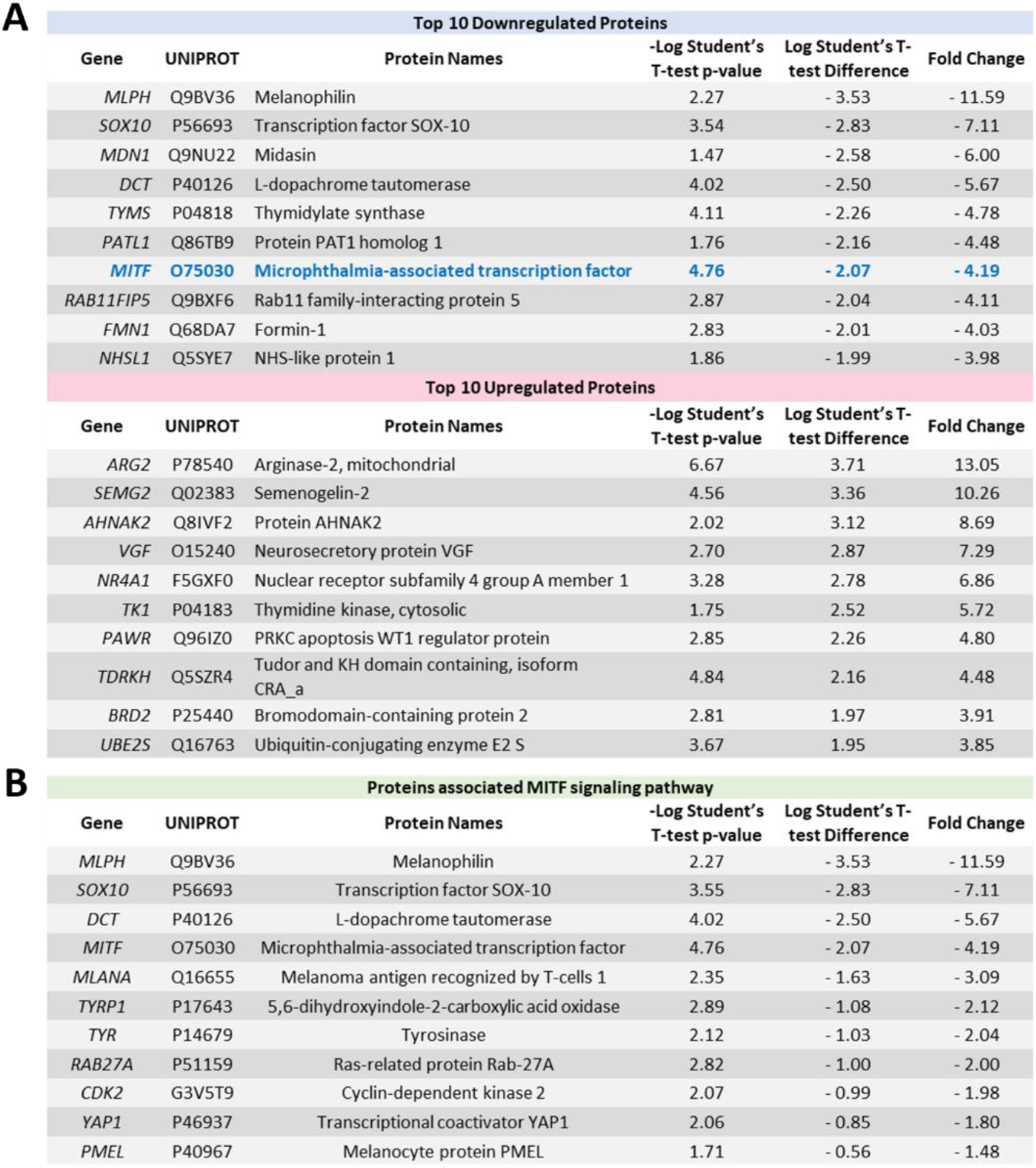
Significantly altered proteins identified by proteomic profiling following ACY-1215 treatment for 24 hours. A: Table highlighting the top 10 strongly down- and up-regulated proteins at 24 hours post treatment (hpt) with 20 μM ACY-1215. B: List of proteins involved in the MITF signaling pathway that were downregulated upon 20 μM ACY-1215 treatment for 24 hours.

Interestingly, from the top 10 downregulated proteins, microphthalmia-associated transcription factor (MITF) was downregulated 4.19-fold by ACY-1215, with proteins connected to MITF signaling also strongly downregulated, *i.e.,* melanophilin (MLPH; 11.59-fold), SRY-box transcription factor (SOX10; 7.11-fold) and L-dopachrome tautomerase (DCT; 5.67-fold), compared to vehicle controls **(Figure 5A)**. Corroborating our proteomics data, MITF expression was significantly downregulated (*p* = 0.002) following 24 hours of 20 µM ACY-1215 treatment **(Figure 6A, 6A’ and Figure S7B)**. A significant difference in MITF expression was not detected after 20 µM ACY-1215 treatment for only 4 hours compared to vehicle control. Expression of additional MITF target proteins and regulators such as Melanoma antigen recognized by T-cells 1 (MLANA; 3.09-fold), 5,6-dihydroxyindole-2-carboxylic acid oxidase (TYRP1; 2.12-fold), Tyrosinase (TYR; 2.04-fold), Ras-related protein Rab-27A (RAB27A; 2.00-fold), Cyclin-dependent kinase 2 (CDK2; 1.98-fold), Transcriptional coactivator YAP1 (YAP1; 1.80-fold), Melanosome protein PMEL (PMEL; 1.48-fold), were significantly reduced by ACY-1215 **(Figure 5B)**. Furthermore, phospho-ERK and ERK expression levels were analyzed in order to determine whether the MAPK/ERK signaling pathway played a role in the ACY-1215 mechanism of action. Through immunoblotting, a significant difference in p-ERK expression levels was not observed after 4 hours of 20 µM ACY-1215 treatment compared to vehicle control **(Figure 6B, 6B’ and Figure S7C)**. Following 24 hpt with 20 µM ACY-1215, p-ERK expression levels were significantly downregulated (*p* = <0.0001) compared to vehicle control **(Figure 6B and Figure S7C)**. Overall, through proteomic analysis, MITF and p-ERK were identified as key players involved in the ACY-1215 mechanism of action in OMM2.5 cells.

**Figure 6:**
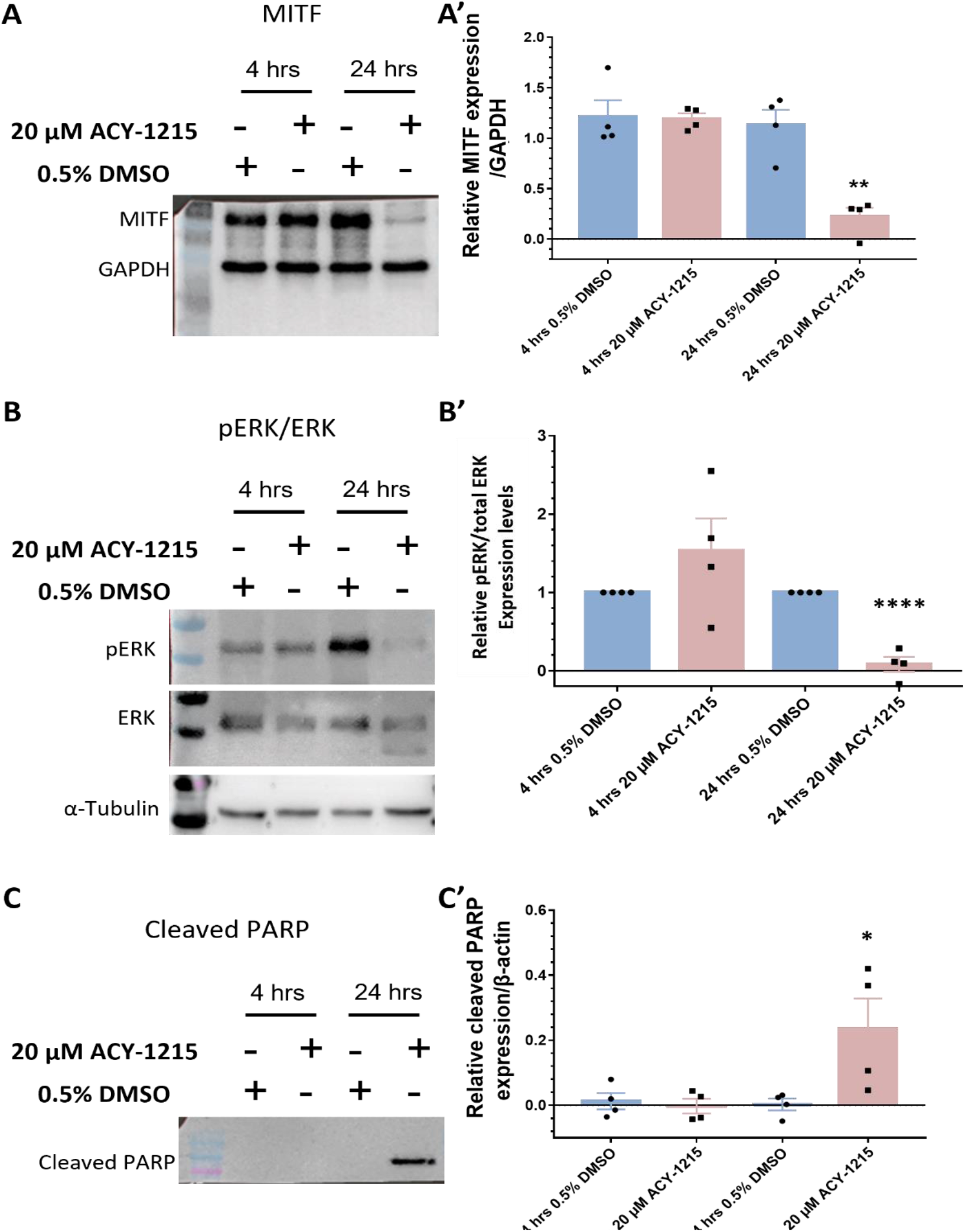
Western Blot validation of proteomics data. A and A’: There was no change in MITF expression levels after 4 hours of 20 μM ACY-1215 treatment, while treatment for 24 hours with 20 μM ACY-1215 led to a significant (**, *p* = 0.002) reduction in MITF expression levels. B and B’: Relative expression levels of p-ERK to total ERK remained unchanged after 20 μM ACY-1215 treatment for 4 hours. At 24 hours post treatment with 20 μM ACY-1215, relative expression levels of p-ERK to total ERK was significantly (****, *p* < 0.0001) downregulated when compared to the 0.5% DMSO treatment. C and C’: Expression of cleaved PARP was significantly (*, *p* = 0.049) upregulated after 24 hours treatment with 20 μM ACY-1215 in comparison to 0.5% DMSO treated OMM2.5 cells. Representative blots for each protein probed and densitometry analysis presented, raw blots are provided in Supplementary Figure 7. β-actin, GAPDH or α- tubulin were used as loading controls. Student’s Unpaired T test statistical analysis was performed, and data presented as mean ± SEM.

### ACY-1215 treatment arrests cell cycle progression in S phase

Outside of UM, previous studies have independently demonstrated that ACY-1215 and MITF regulate the cell cycle [40–44]. To determine whether ACY-1215 treatment altered cell cycle phases in MUM cells, OMM2.5 cells were treated with either 0.5% DMSO, 10, 20 or 50 µM of ACY-1215, 50 µM Etoposide (a chemotherapeutic used as a positive control for apoptotic cell death) or 20 µM Dacarbazine for 4 and 24 hours. The cells were isolated, fixed, labelled with propidium iodide, and analyzed using flow cytometry **(Figure 7 and Figure S8)**. In line with published studies, OMM2.5 cells undergo two cell cycle phases, due to the DNA ploidy of UM cells [45, 46]. Approximately 60% - 70% of the cell population were diploid, in cell cycle 1 and the remaining cell population presented with aneuploidy in cell cycle 2 **(Figure S8)**. Significant changes in G1, S and G2 cell cycle phases were not observed after 4 hours of ACY-1215 in any treatment group compared to vehicle controls **(Figure 7B, 7C and 7E)**. After 24 hours of treatment with Etoposide or ACY-1215, a significant reduction (*p* = 0.0001) in the number of cells in G1 phase and a significant increase (*p* = 0.0001) in the number of cells in S phase was identified across the treatment groups compared to vehicle controls **(Figure 7B, 7D and 7F)**. On average, 19.0%, 8.2% and 11.9% of OMM2.5 cells were in G1 phase following 10, 20 and 50 µM of ACY-1215 treatment, respectively, compared to 58.0% of cells in G1 when treated with vehicle control. 80.7%, 91.6% and 88.0% of cells on average were detected in S phase upon treatment with 10, 20 and 50 µM of ACY-1215 in comparison to 39.35% of 0.5% DMSO treated cells. OMM2.5 cells treated with 20 µM Etoposide had 7.8% of cells in G1 and 88.4% of cells in S phase. The number of cells in G2 phase across all treatment groups did not significantly change at 24 hpt. No change was observed in any of the cell cycle phases following Dacarbazine treatment at 4 or 24 hours. In summary, cell cycle analysis proved that ACY-1215 treatment for 24 hours attenuated OMM2.5 cell cycle progression in S phase.

**Figure 7:**
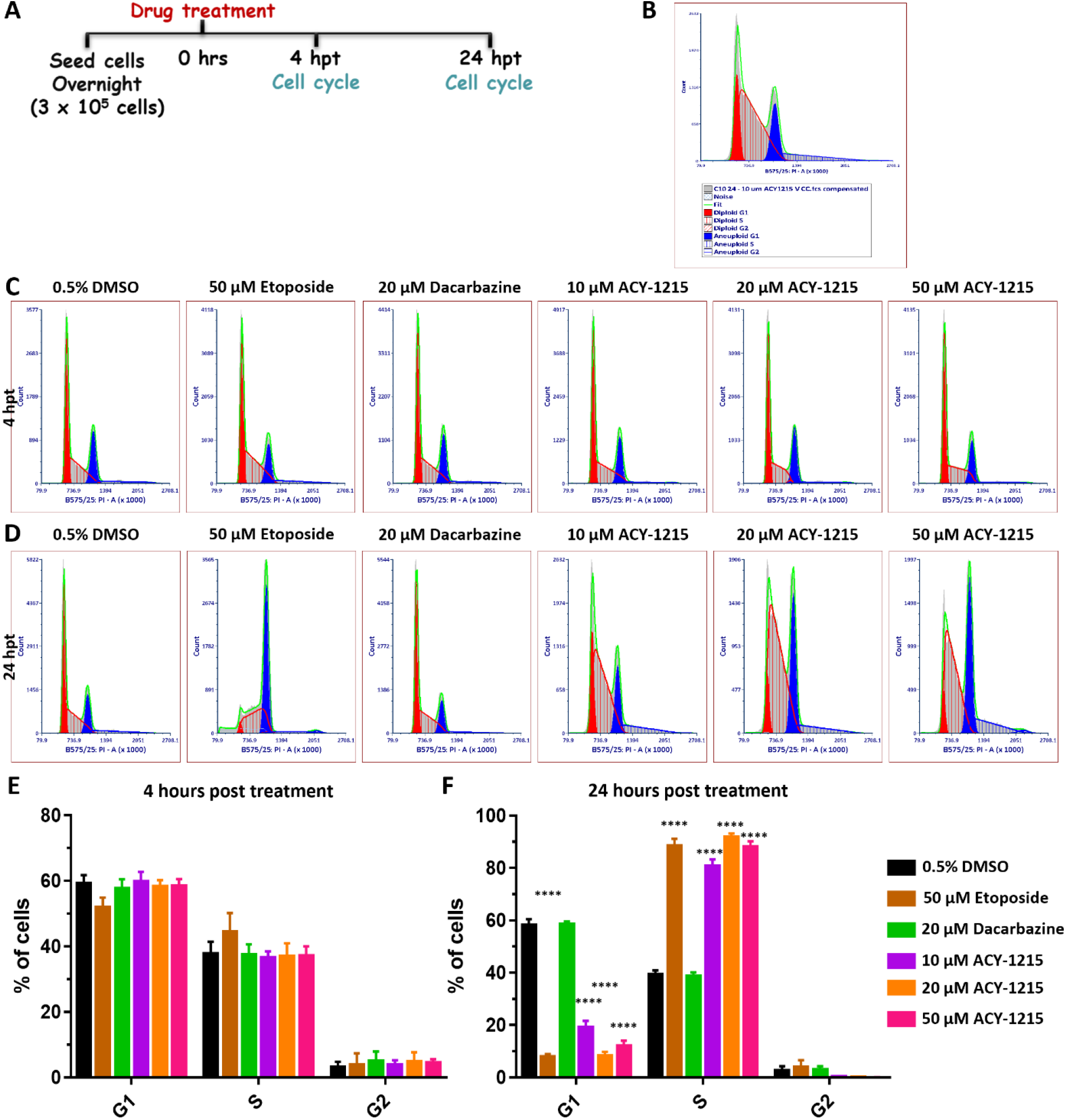
Cell cycle progression is arrested in S phase by ACY-1215 in OMM2.5 cells. A: Schematic illustrating treatment protocol undertaken. B: Flow cytometry data analysis plot legend. C and E: 4 hours of treatment with 10, 20 or 50 μM ACY-1215; or 50 μM Etoposide or 20 μM Dacarbazine did not alter the cell cycle profile. D and F: A significant (****, *p* = 0.0001) reduction in the percentage of cells in G1 phase and a significant (****, *p* = 0.0001) increase in the percentage of cells in S phase was observed after 24 hpt with 10, 20 or 50 μM ACY-1215 or 50 μM Etoposide, in comparison to vehicle controls. No changes in the cell cycle phases were observed following 20 μM Dacarbazine treatment. No alterations to G2 phase were observed in all treatment groups. Statistical analysis was performed by two-way ANOVA with Dunnett’s Test for Multiple Comparisons and data represented as mean ± SEM (N = 4).

### Elevated apoptosis results from ACY-1215 treatment of MUM cells

As the majority of OMM2.5 cells were arrested at S phase after 24 hours of ACY-1215 treatment, we investigated if these cells undergo increased apoptosis. OMM2.5 cells were treated, isolated, labelled with YO-PRO^TM^-1 Iodide and Propidium iodide to distinguish between viable, non-viable and cells in different apoptotic stages **(Figure 8 and Figure S9)**. In line with our cell cycle results, 4 hours of ACY-1215 treatment did not significantly alter apoptotic cell number in any treatment group **(Figure S9)**. At 24 hpt, a significant reduction in live cells was reported with 20 µM (2.52% reduction of total number of live cells; *p* = 0.0055) and 50 µM (5.28% reduction of total number of live cells; *p* = <0.0001) ACY-1215 compared to the vehicle control **(Figure 8A’, 8A’’ and 8C)**. Additionally, ACY-1215 significantly increased the average number of early apoptotic cells as evidenced by 3.22% (*p* = 0.017) and 4.89% (*p* = <0.0001) early apoptotic cells following 20 µM or 50 µM ACY-1215 treatment, respectively, compared to the vehicle control. After 24 hours of treatment, there was no significant difference detected in the average number of cells undergoing late apoptosis or dead cells across all treatment groups **(Figure 8A’’ and 8C)**. In line with our findings, cleaved PARP expression (a marker for apoptosis) was significantly upregulated at 24 hpt with 20 µM ACY-1215 (*p* = 0.049) and not at 4 hpt **(Figure 6C, 6C’ and Figure S7D)**.

**Figure 8:**
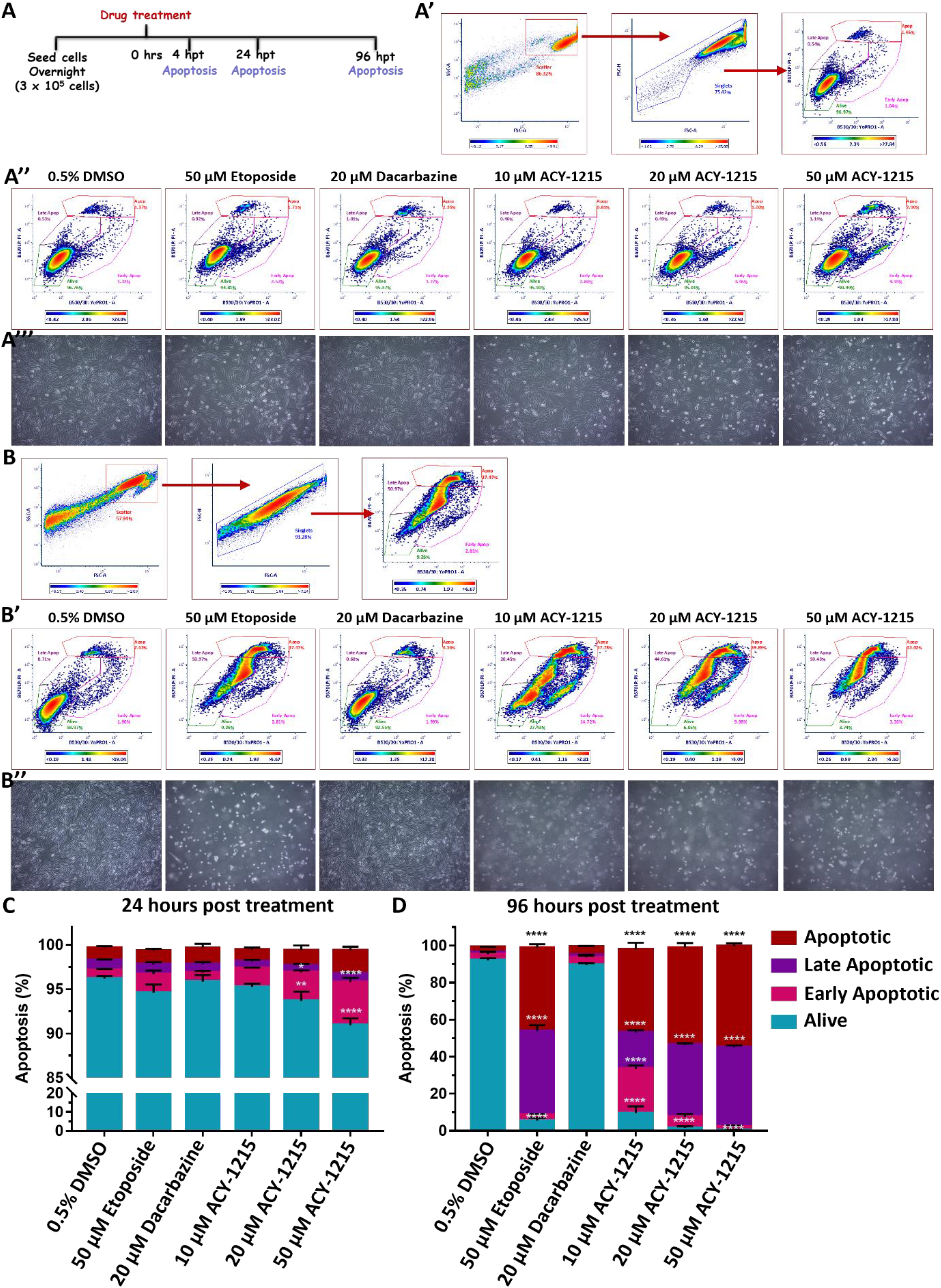
ACY-1215 activates the apoptotic pathway in OMM2.5 cells. A: Diagram portraying treatment regime. A’ and B: Plots representing gating of cell singlets into different stages of apoptosis. A’’ and B’: Representative plots depicting OMM2.5 treated with 0.5% DMSO; 10, 20 or 50 μM ACY-1215; 50 μM Etoposide or 20 μM Dacarbazine, at 24 and 96 hpt, respectively. A’’’ and B’’: Representative micrographs of OMM2.5 cells at 24 and 96 hours post treatment. C: A significant reduction in the percentage of live cells and a significant increase in the percentage of early apoptotic cells was detected following 20 μM (**, *p* = 0.005 and *, *p* = 0.016, respectively) and 50 μM (****, *p* < 0.0001) ACY-1215 treatment compared to 0.5% DMSO treatment. D: Live cell populations were significantly (****, *p* < 0.0001) reduced and cell populations in late apoptotic stage and apoptotic stage were significantly (****, *p* < 0.0001) increased upon 50 μM Etoposide and all concentrations of ACY-1215 tested. 10 μM ACY-1215 treatment resulted in a significant increase in the early apoptotic cell population compared to 0.5% DMSO. 20 μM Dacarbazine treatment was comparable to vehicle control plots. Statistical analysis by Two-way ANOVA followed by Tukey’s Multiple Comparisons test with error bars shown as mean ± SEM (N = 3).

Prolonged ACY-1215 treatment for 96 hours, resulted in the majority of cells being either non-viable or undergoing late apoptosis **(Figure 8B, 8B’ and 8D)**. The average number of viable cells with 10, 20 or 50 µM ACY-1215 was significantly reduced to 9.47% (*p* = <0.0001), 1.56% (*p* = <0.0001) and 0.46% (*p* = <0.0001), respectively. In contrast 92.9% and 89.45% of cells were viable in vehicle control and 20 µM Dacarbazine treated groups, on average, respectively **(Figure 8B’ and 8D)**. A significant increase in early apoptotic cells was detected in the 10 µM ACY-1215 treatment with 23.95% (*p* = <0.0001) of cells, compared to the vehicle control; a significant change was not observed in all other treatment groups. In 0.5% DMSO treatment group 1.15% of cells and 1.36% of cells treated with 20 µM Dacarbazine were in late apoptotic stage while a substantial number of cells, on average 19.8% (*p* = <0.0001) in 10 µM, 39.0% (*p* = <0.0001) in 20 µM and 42.9% (*p* = <0.0001) in 50 µM ACY-1215 treated groups were undergoing late-stage apoptosis **(Figure 8B’ and 8D)**. ACY-1215 treatment resulted in a profound number of non-viable cells in a dose-dependent manner, with 44.8% (*p* = <0.0001), 52.4% (*p* = <0.0001) and 54.5% (*p* = <0.0001) following 10, 20 and 50 µM concentrations, in comparison to 2.71% dead cells in vehicle control and 4.47% in 20 µM Dacarbazine treated groups **(Figure 8B’ and 8D)**. Etoposide (50 µM), a positive control for apoptosis, showed 5.41% (*p* = <0.0001) cells were viable, 45.2% (*p* = <0.0001) were in late apoptotic stage and 45.0% (*p* = <0.0001) were non-viable **(Figure 8B’ and 8D)**. Furthermore, micrograph images of all treated cells corroborate our results that 96 hours of treatment with Etoposide or ACY-1215 significantly reduced cell viability, with most of the cells not adhered to the culture plate, in contrast to the vehicle control or clinical chemotherapeutic for 24-hour treatment groups **(Figure 8A’’’ and 8B’’)**. Overall, we observe a time- and dose dependent alteration in OMM2.5 cell viability, cell cycle arrest and triggering of apoptosis, 24 hours post ACY-1215 treatment.

### MITF inhibitor treatment prevents MUM cell survival and proliferation in vitro

To further interrogate the requirement of MITF in MUM cell survival, the ability of OMM2.5 cells treated with the MITF pathway inhibitor ML329, to survive and proliferate was analyzed using colony formation assays. Cells were treated with increasing doses of ML329, ranging between 0.05 µM and 50 µM, given the reported IC50 value of 1.2 µM **(Figure 9A and 9B)** [47]. The treatment regime was performed aspreviously described, whereby OMM2.5 cells were treated with respective drug doses for 96 hours, and then maintained in culture, in fresh complete media for an additional 10 days **(Figure 9A)**.

**Figure 9:**
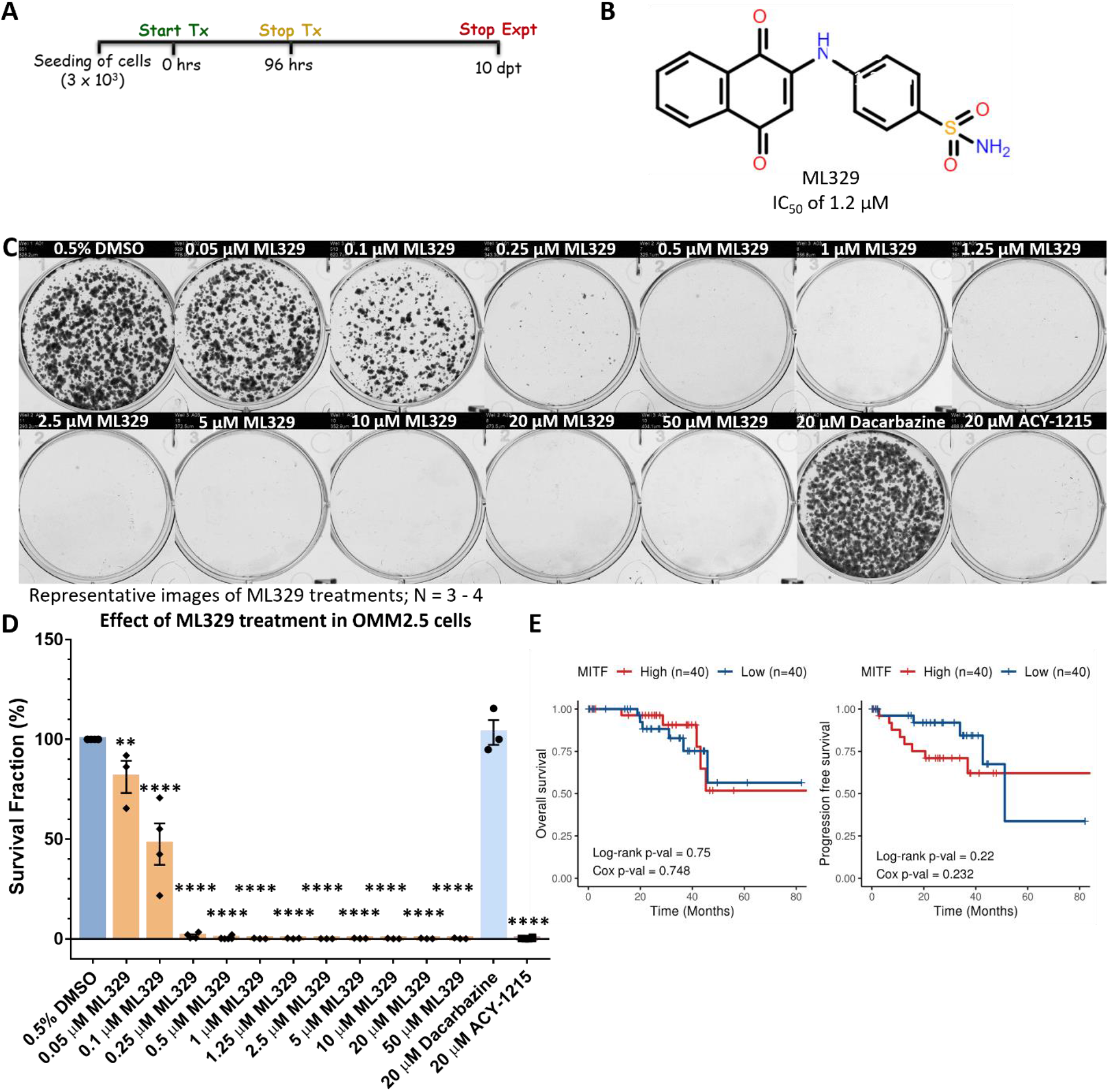
Inhibition of MITF pathway reduces OMM2.5 cell viability *in vitro*. A: Schematic diagram of treatment regime. B: Chemical structure of ML329, a small molecule MITF pathway inhibitor. C: Representative image of clonogenic assay plates for OMM2.5 cells treated with 0.5% DMSO, 0.05 - 50 μM ML329, 20 μM Dacarbazine or 20 μM ACY-1215 for 96 hours. D: A dose-dependent, significant reduction in the number of OMM2.5 colonies was observed in ML329 treatment groups compared to the 0.5% DMSO treatment group. One-way ANOVA with Dunnett’s Test for Multiple Comparisons statistical analysis was performed, error bars represent mean ± SEM, ****, *p* value of 0.0001 (N = 3 - 4). E: Kaplan-Meier survival curves demonstrating no correlation between expression of ML329 and overall survival (OS) or progression free survival (PFS) in UM patients. Median values were used as cut-off for high (red) and low (blue) expression levels, with Log-rank p-values (categorical variable) and Cox p-values (continuous variable) calculated (n = 80).

ML329 induced a significant reduction in the average number of surviving colonies (reduced by 18.9%, *p* = 0.005) when treated with 0.05 µM ML329 treatment compared to 0.5% DMSO **(Figure 9C and 9D)**. At higher concentrations of ML329, more pronounced effects were detected with significant reductions in viable clones averaging 52.6% to 99.8% (*p* = 0.0001) decreases at 0.1 to 50 µM concentration of ML329, compared to vehicle controls **(Figure 9C and 9D)**. Corroborating our data, treatment of OMM2.5 cells with 20 µM Dacarbazine did not result in a significant difference in the average number of viable clones while 20 µM ACY-1215 treatment led to a significant reduction (99.8%; *p* = 0.0001) in number of surviving clones, compared to 0.5% DMSO. Given that MITF was found to play a role in MUM cell survival, correlations between *MITF* expression and UM patient OS/PFS was investigated. Curiously, high or low *MITF* expression levels were not significantly associated with better OS nor PFS (Cox OS, *p* = 0.748 and Cox PFS, *p* = 0.232) as shown by Kaplan-Meier survival curves **(Figure 9E)**.

### Inhibition of MITF pathway hinder MUM cells survival in vivo in zebrafish OMM2.5 xenograft models

The efficacy of the MITF pathway inhibitor, ML329, on survival of MUM cells *in vivo* was determined using zebrafish xenograft models. ML329 was well tolerated by zebrafish *in vivo*, albeit with drug precipitation at higher concentrations (1 - 100 µM) **(Figure S10).** Although we observed effects *in vitro* at concentrations as low as at 0.25 µM ML329, we chose the concentration of 1.25 µM for our study to fit with the reported IC50 value [47]. As before, OMM2.5 Dil labelled cells were injected into the perivitelline space after which the larvae (2 dpf) were treated with either 0.5% DMSO or 1.25 µM ML329 for 3 days **(Figure 10A)**. There was no significant difference in the average number of disseminated cells to the caudal vein plexus of the OMM2.5 xenografted larvae at 0.5% DMSO (3.1 cells) or 1.25 µM ML329 (2.6 cells) treatment groups **(Figure 10B and 10D)**. However, on average, a 51% (*p* = 0.0006) reduction in OMM2.5 primary xenograft fluorescence was detected after normalization, following treatment with 1.25 µM ML329 compared to vehicle controls **(Figure 10A and 10C)**. Experimentally, therefore we observe a beneficial effect of blocking the MITF pathway in MUM cell line *in vitro* and *in vivo*.

**Figure 10:**
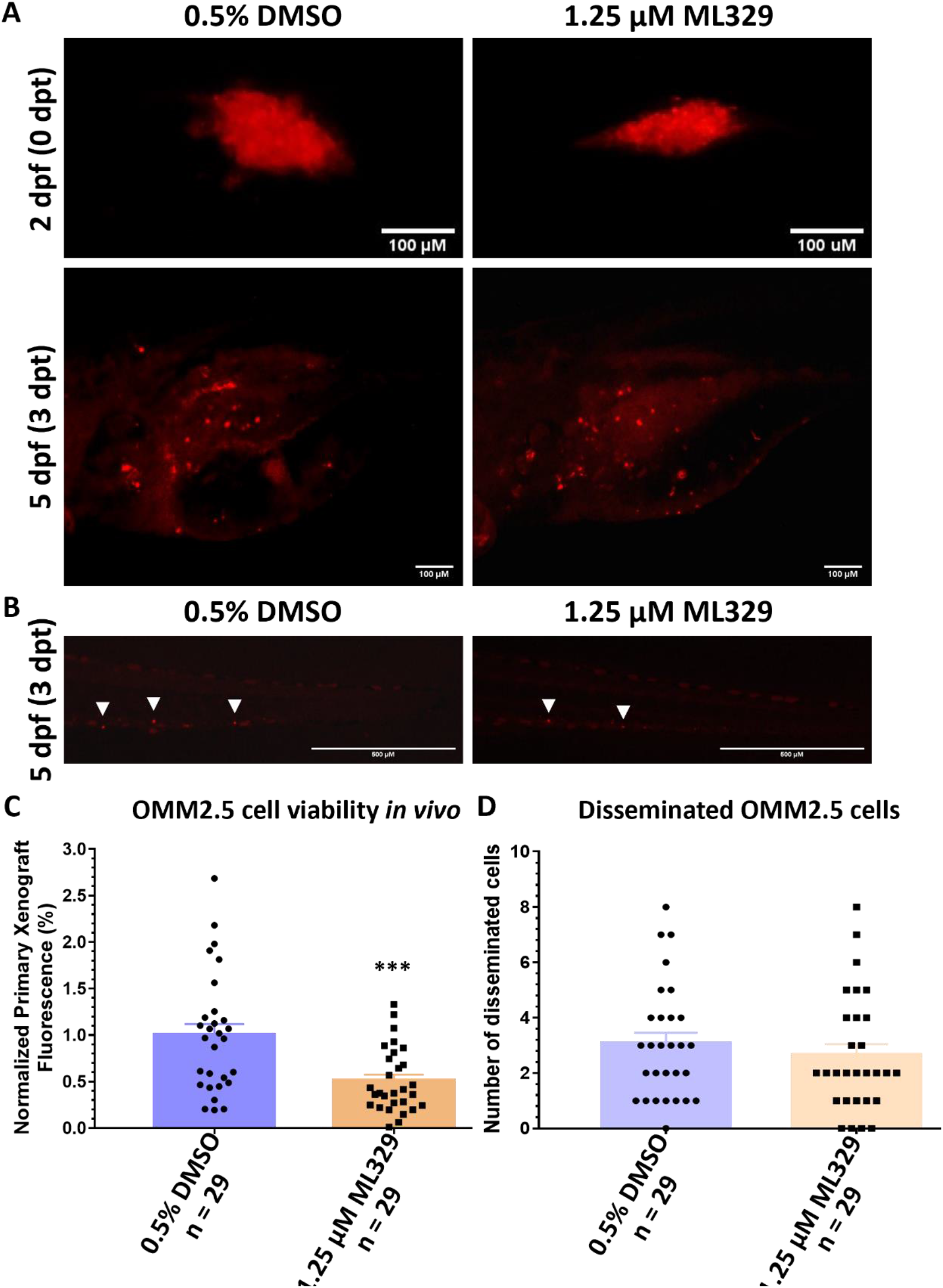
ML329 demonstrates anti-UM properties in zebrafish OMM2.5 xenografts. A: Top panel shows OMM2.5 Dil labelled cells xenografted into the perivitelline space of 2 days old zebrafish larvae. Bottom panel presents zebrafish larvae 3 days post treatment with 0.5% DMSO (n = 29) or 1.25 μM ML329 (n = 29). B. Representative image of OMM2.5 Dil labelled cells disseminated (white arrowhead) to the caudal vein plexus of zebrafish larvae at 3 dpt. C: A significant (***, *p* = 0.0006) regression of the average normalized primary xenograft fluorescence of OMM2.5 Dil labelled cells was observed when treated with 1.25 μM ML329. D: No difference detected in the average number of disseminated OMM2.5 Dil cells following treatment with 1.25 μM ML329 compared to vehicle control. Student’s T test was used for statistical analysis with error bars presenting mean ± SEM.

## Discussion

Metastatic UM (MUM) is a poor prognosis cancer lacking effective treatment options. Our study has provided evidence that inhibition of HDAC6 or MITF is efficacious in conferring anti-cancer effects in a MUM cell line, both *in vitro* and *in vivo*. To the best of our knowledge, this is the first study to provide evidence regarding the potential link between HDAC6 and MITF in MUM.

Three commercially available, first generation HDAC6i were screened in UM and MUM cell lines and ACY- 1215 was selected for follow-up studies. ACY-1215 either as a monotherapy or in combination with other drugs, is presently in clinical trials for several cancers [27, 48]. We observed strong anti-cancer effects elicited by ACY-1215 treatment in a dose-dependent manner in both UM and MUM derived cell lines, albeit weak HDAC6 expression is reported in UM tissues [49]. Notably, HDAC6 activity is significantly increased in inflammatory breast cancer even though HDAC6 is not overexpressed [50]. Hence, it is plausible that in MUM, there is increased HDAC6 activity but not HDAC6 expression. Our data indirectly supports the findings by Nencetti *et al.*, whereby a novel synthetized quinoline derivative VS13, with high selectivity against HDAC6; led to a reduction in UM cell viability *in vitro* [26]. In addition, here, the anti- cancer effect of ACY-1215 was demonstrated *in vivo* in zebrafish OMM2.5 xenograft models, without any significant change to the number of disseminated cells. This is not surprising given the timeframe of the experiment, and a low burden in the average number of disseminated cells to the caudal vein plexus three days post transplantation in the vehicle controls. It would be worthwhile to perform follow-up studies to evaluate the efficacy of ACY-1215 on tumor metastasis, with long-term treatment regimens and in patient-derived samples *in vivo* in larvae and/or in adult zebrafish [51–54].

However, pure HDAC6 inhibition mediated effects must be inferred with caution, as higher doses of ACY- 1215, result in non-selective inhibition, and the observed beneficial effects are mediated by additional targets [39, 55]. In a study by Lin *et al.,* CRISPR-induced HDAC6 knock-out lines (*e.g.* melanoma, triple negative breast cancer, colorectal cell lines) demonstrated that the cell viability/proliferation capability was comparable to wildtype controls; additionally ACY-1215 was able to mediate its anti-cancer effects at high concentrations (micromolar) even when HDAC6 was knocked-out [55]. Corroborating their findings, Depetter *et al.*, revealed that treatment with 10 μM ACY-1215 in HAP1 cells with HDAC6 knock-out, led to a reduction in cell viability [39]. In another study, a distinct anti-proliferative effect was observed in high-grade serous ovarian cancer cells when a non-selective concentration of 10 μM ACY-1215 was used [56]. In both studies, the authors suggest that the true beneficial effects of HDAC6 inhibition might be reaped in combinatorial therapy rather than when administered as a single agent. Therefore, it has to be acknowledged that at our selected treatment concentration of 20 μM, we are likely to be non-selectively targeting other factors such as Class I HDAC isozymes, given the reported IC50 value for ACY-1215 is 4.7 nM. Importantly, HDAC6 was indeed inhibited by ACY-1215 at the concentration we used, as its substrate acetylated α-tubulin was significantly upregulated. Furthermore, from our proteomics data we also identified proteins involved in *microtubule polymerization* and *regulation of microtubule polymerization or depolymerization* to be significantly altered [57]. Irrespective of non-selective inhibition of HDAC isozymes, ACY-1215 still presents as a promising therapeutic for treatment of MUM, with its ability to prevent UM cell growth, that warrants further interrogation.

Proteome profiling of ACY-1215 treated OMM2.5 cells was key to deducing potential mechanisms of action. We discovered that the MITF signaling pathway and associated factors were significantly downregulated upon treatment with ACY-1215. Tying in with the concentration of ACY-1215 used, our findings are in line with another study, whereby it was reported that treatment of melanoma and clear cell sarcoma cells with different pan-HDAC inhibitors resulted in reduced MITF expression *in vitro* and *in vivo* in a mouse melanoma xenograft model [58].

The role of MITF has been extensively studied in cutaneous melanoma [59–61]. MITF is a key transcription factor and a master regulator of melanogenesis and melanocyte differentiation. It also plays a multifaceted role regulating several cellular processes including cell cycle, DNA-damage repair, lysosome biogenesis, metabolism, autophagy, and oxidative stress [62–65]. MITF can be further distinguished into five different isoforms, MITF-A, MITF-B, MITF-C, MITF-H and MITF-M [66]. Particularly in cutaneous melanoma, MITF-M is involved in carcinogenesis events such as survival, proliferation, differentiation, invasion and migration [61]. Not surprising, certain types of mutations in MITF and MITF-associated members are linked to oncogenic functions in melanoma [62,67,68]. MITF plays a dual role in cutaneous melanoma, based on its expression levels and activity, however, there is controversy surrounding this matter [63]. For instance, some studies report that low MITF expression is necessary for proliferation and higher levels of MITF correlates to suppression of cell proliferation and promotes differentiation [61]. While others state that low levels of MITF expression is linked to invasiveness while high levels of MITF expression is required for cell proliferation/differentiation [42,60,69]. Nevertheless, targeting the MITF pathway shows promise as an anti-cancer approach. Aida *et al.*, demonstrated that the growth of melanoma cells, SK-MEL-5 and SK-MEL-30 were inhibited by siRNA mediated knock-down of MITF [70]. Similarly, in another study, knock-down of MITF by shRNA, in MM649 cells resulted in reduced cell proliferation *in vitro* and tumor growth and dissemination *in vivo* in mouse xenografts [59]. Furthermore, pharmacological inhibition of the MITF signaling pathway using small molecule ML329 reduced cell viability in MITF-dependent melanoma (SK-MEL-5 and MALME-3M) cells without affecting the viability of A375 cells, a MITF-independent cell line [47]. Comparably, another compound, CH5552074, inhibited the growth of SK-MEL-5 cells via the suppression of MITF protein [70]. Interestingly, knock-down of MITF in B16F10 melanoma cells and overexpression of MITF in YUMM1.1 cells led to increased tumor growth *in vivo* in mice [71]. Apart from melanoma, studies have connected MITF with a role in multiple cancers including non-small cell lung cancer, pancreatic cancer, and hepatocellular carcinoma [72–74]. Most recently, it was demonstrated that knockdown of MITF in clear cell renal cell carcinoma cells, resulted in reduced cell proliferation and an increase in cells in S/G2 phases, suppressed cell migration and invasion *in vitro* and tumor formation *in vivo*; an opposing effect was observed when MITF was overexpressed [43]. In the context of UM, MITF is upregulated in UM cells [75]. In our study, expression levels of MITF and several proteins involved in pathways associated with MITF, such as pigment cell differentiation and melanosome organization, were downregulated upon ACY-1215 treatment. This was consistent with the observed trend when MITF is downregulated. Taken together, there is ample evidence to suggest that targeting the MITF signaling pathway may be a novel therapeutic option for MUM.

Moreover, several studies have independently shown that ACY-1215 regulates cell cycle and cell death mechanisms in various cancers. In HCT-116 and HT29 colorectal cancer cells, a reduction in cell proliferation and viability was noted in a time- and dose-dependent manner; and apoptosis was observed as well at non-selective ACY-1215 concentrations [41, 76]. Interestingly, ACY-1215 when used at HDAC6 selective concentrations (up to 2 μM) did not promote apoptosis, however, if used in combination with other anti-cancer drugs it proved to be more effective [76, 77]. In esophageal squamous cell carcinoma cell lines (EC109 and TE-1), ACY-1215 treatment resulted in suppression of cell proliferation through the arrest of cell cycle in G2/M phase and an increase in apoptosis [78]. Similarly, 4 μM ACY-1215 treatment for 24 hours, prompted an increase in percentage of cells in G0/G1 phase; and a time/dose-dependent proapoptotic effects of ACY-1215 uncovered in lymphoma cell lines [40]. More recently, in gall bladder cancer cells, ACY-1215 inhibited cell proliferation and induced apoptosis as well as enhancing the chemotherapeutic effects of other anti-cancer agents upon co-treatment [79]. Collectively, in these studies it became evident that the PI3K/AKT and MAPK/ERK pathways played a central role in ACY-1215 mechanism of action. We postulated whether ACY-1215 treatment promoted cell cycle arrest and apoptosis in MUM cells. As expected, at the non-selective concentration, ACY-1215 treatment resulted in the halting of cell cycle progression in S phase and induced apoptosis. We observed a significant increase in early apoptotic cells and a significant reduction in the number of viable cells at 20 and 50 μM ACY-1215 treatment by 24 hours. Additionally, expression of cleaved PARP, which is used as an indicator for apoptosis, was markedly upregulated in ACY-1215 treated MUM cells at 24 hours post treatment [80, 81]. Further supporting evidence can be drawn from our proteomics data, whereby the pathways - *regulation of extrinsic apoptotic signaling pathway in absence of ligand*, *exit from mitosis* and *cellular senescence* were upregulated indicating an increase in expression levels of proteins associated with these biological processes. By 96 hours, at all tested ACY-1215 concentrations, the majority of cells were either apoptotic or in late apoptotic stages. Considering that MITF was significantly downregulated at 24 hours post treatment with ACY-1215, the cause of increased cell death observed following ACY-1215 treatment, is potentially mediated through the downregulation of MITF. In order to further confirm that the observed anti-cancer effects of ACY-1215 is through the regulation of MITF, OMM2.5 cells were treated with a MITF pathway inhibitor, ML329, *in vitro* and *in vivo* in zebrafish OMM2.5 xenografts. We noted a dose-dependent reduction in cell viability *in vitro* and at the tested concentration, inhibition of the MITF pathway revealed an anti-UM effect *in vivo*. Additional interrogation of the link between HDAC6 and MITF, opened up the possibility of MAPK/ERK signaling being involved in the ACY-1215 mechanism of action in MUM cells. We observed that p-ERK expression levels were significantly reduced following 24 hours of ACY-1215 treatment. This correlates with reduced MITF expression levels, observed at the same time. It is not surprising that the MAPK/ERK signaling pathway is involved in ACY-1215 mechanism of action. It was previously reported that ERK1/2 and HDAC6 are interacting partners involved in a positive feed- forward loop [82–84]. In colon cancer cell lines, knock-down of HDAC6 resulted in reduced p-ERK expression but not total ERK expression levels [85]. Peng *et al*., showed that in A375 melanoma cells, inhibition with ACY-1215 alone and in combination with vemurafenib led to the disruption and inactivation of ERK [86]. Interestingly in prostate cancer cells (LNCaP), blocking of HDAC6 with Panobinostat led to increased ERK activity and as a consequence promoted apoptosis [87]. But this was not the case in PC-3 prostate cancer cells. While in another study, increased HDAC6 expression in lung cancer cell by Isoproterenol treatment led to the inhibition of the ERK signaling cascade [88]. Taken together, this indicates that there might be cell-specific context for HDAC6-ERK1 regulation and activity. In UM, *GNAQ/GNA11* mutations are associated with constitutive activation of the MAPK/ERK signaling pathway, although heterogeneity in MAPK/ERK signaling has been observed across UM samples with *GNAQ/GNA11* mutations [89–92]. More specifically, the OMM2.5 cells used in this study carry a mutation in *GNAQ*, which is known to result in constitutively active MAPK/ERK signaling in UM [8, 93]. Therefore, we postulate that inhibition of HDAC6 leads to reduced ERK activity that subsequently results in reduced p- ERK expression levels. In turn, reduced p-ERK expression levels decrease MITF expression and consequently, associated biological processes downstream such as cell survival mechanisms are inhibited **(Figure 11).** Collectively, our findings warrant an in-depth analysis into understanding the role of MITF in MUM and to consider targeting MITF and/or candidates in the MITF signaling pathway as additional potential therapeutic option(s).

**Figure 11:**
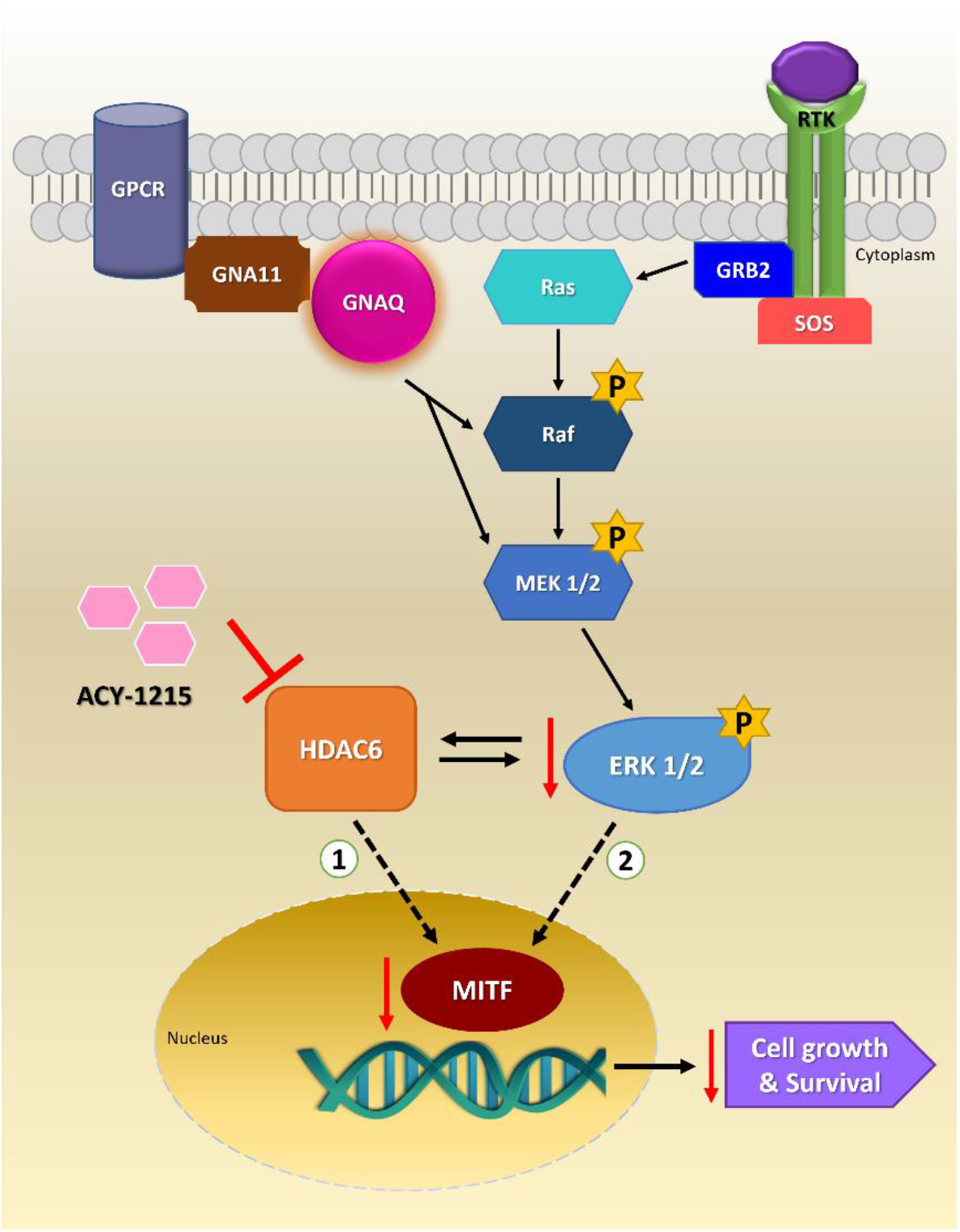
Proposed model for ACY-1215 mechanism of action in OMM2.5 cells. Mutations in GNAQ, lead to constitutively active MAPK/ERK signaling that activates cell survival mechanisms in OMM2.5 cells (black arrows). Inhibition of HDAC6 by ACY-1215 is either ① directly involved in the downregulation of MITF expression levels and consequently its downstream effectors and biological processes; or ② downregulation of p-ERK expression levels upon HDAC6 inhibition, in turn downregulates MITF expression and consequently inhibits biological processes that promotes cell survival and proliferation. Red arrows represent consequence of HDAC6 inhibition and dotted arrows indicative of plausible mechanism of action.

Though promising, the role of HDAC6 and MITF still needs to be thoroughly investigated in UM and MUM patient samples. Currently, there is no clear evidence linking either HDAC6 or MITF in MUM prognosis. Immunohistochemistry based expression analysis of 16 primary UM samples detected variable low levels of HDAC6 expression, with a clear correlation between HDAC6 expression levels and UM, unable to be drawn due to the limited sample size [49]. Based on TCGA data analysis comprising 80 UM patient samples, a significant correlation was found between *HDAC6* expression and OS probability, highlighting a possible involvement of HDAC6 in UM prognosis. Moreover, *HDAC2* and *SIRT2* expression correlated to OS while *HDAC4* expression showed correlations to PFS. HDAC 1 and 3 expression was not correlated to either OS or PFS. Although, HDAC 1, 2, 3, 4 and Sirtuin 2 (SIRT2) expression was detected in UM eye samples [49]. A limited number of studies have explored the expression of MITF in UM and MUM. MITF expression was found in 100% (15 out of 15) of 15) of UM samples in one study, however, in another study, MITF expression was detected in 65% (37 out of 57 samples) of choroidal UM patient samples, with levels of MITF expression not significantly associated with the survival of these patients [94,95]. Comparably, from our TCGA data analysis, there was no correlation between MITF expression levels and OS/PFS seen in UM patients. It has been previously suggested that MITF would be a useful marker for ocular malignant melanoma [96]. Taken together, it will be worthwhile to perform an extensive study with a larger cohort of UM and MUM patient samples to conclusively determine whether HDAC6 and MITF plays a part in MUM prognosis. Additionally, it needs to be determined whether targeting these pathways offer a broad treatment option for MUM irrespective of MUM causative mutation(s).

This study is pivotal in highlighting that MITF plays a critical role in the survival of OMM2.5 cells and provides evidence that the observed ACY-1215 mechanism of action in OMM2.5 cells is most likely through the regulation of MITF. Our data suggests that HDAC6 and/or pan-HDACs and the MITF signaling pathway offer novel options to identify therapeutic targets for treatment of MUM that needs to be considered and further evaluated.

## Methods

### Cell culture

Mel270, Mel285 and OMM2.5 cells (kindly provided by Dr. Martine Jager, Leiden, The Netherlands/Dr. B.R. Ksander, Schepens Eye Research Institute, Boston, USA) were cultured with Complete Media containing RPMI 1640 (Gibco; Waltham, MA, United States) + 10% fetal bovine serum (FBS) + 2% penicillin-streptomycin (PEST), in T175 flasks for not more than 20 passages [35]. ARPE-19 cells were maintained in Complete media containing DMEM: F12 supplemented (Lonza; Basel, Switzerland) + 10% FBS + 1% PEST + 2.5 mM L-Glutamine. Culture flasks and plates were incubated at 37^0^C with 5% CO_2_.

### Clonogenic assay

All three cell lines were seeded into 6-well plates at 1.5 x 10^3^ cells/ml (final volume 2 ml) and allowed to adhere overnight. Initial drug screens were performed with Mel285 cells seeded at 1.5 x 10^3^ cells/ml and Mel270/OMM2.5 cells were seeded at 9 x 10^3^ cells/ml. The following day, cells were treated with either 0.5% DMSO (vehicle control) or 20 µM Dacarbazine (clinical control; Sigma-Aldrich; St. Louis, MO, United States) or HDAC6i (Tubastatin A (SelleckChem; Houston, TX, USA), ACY-1215 (SelleckChem) and Tubacin (Sigma-Aldrich)) at increasing doses ranging from 1 µM, 5 µM, 10 µM, 20 µM and 50 µM, prepared in Complete Media. The MITF inhibitor, ML329 (Ambeed, Inc.; Sigma-Aldrich), was tested at increasing concentrations ranging from 0.05 µM, 0.1 µM, 0.25 µM, 0.5 µM, 1 µM, 1.25 µM, 2.5 µM, 5 µM, 10 µM, 20 µM and 50 µM. All drugs were dissolved in DMSO to prepare stock solutions. Cells were treated with 2 ml of desired drug solution per well in duplicate and incubated at 37^0^C with 5% CO_2_ for 96 hrs. Drug solutions were removed and wells washed twice with 1 x phosphate buffered saline (PBS; Lonza). Fresh Complete

Media were added to the plates and cells allowed to grow for an additional 10 days at 37^0^C with 5% CO_2_. Clones were washed twice and fixed with 4% paraformaldehyde/formaldehyde for 10 mins at room temperature (RT). Clones were stained with 0.5% crystal violet solution (Pro-Lab diagnostics PL700; Richmond Hill, ON, Canada) for 10 mins - 2 hours at RT, shaker, washed and dried (once desired staining is achieved). Plates were imaged using the GelCount™ system (Oxford Optronix; Oxford, UK) and analyzed using the ColonyCountJ Plugin in ImageJ v1.53e [97]. Statistical analysis was performed using One-way ANOVA with Dunnett’s Test for Multiple Comparisons in GraphPad Prism v7.00 for Windows (GraphPad Software, San Diego, CA, USA, www.graphpad.com). A *p* value of ≤ 0.05 was considered as statistically significant. Experiments were performed in triplicates/quadruplicates.

### OMM2.5 zebrafish xenografts

All animal work and husbandry were performed in accordance with ethical approval granted by Linköping Animal Research Ethics Committee. Zebrafish embryos/larvae from *Tg(fli1a:EGFP)* background were raised in embryo media containing 5 mM NaCl, 0.17 mM KCl, 0.33 mM MgCl2, 0.33 mM CaCl2 and 0.003% phenylthiourea (PTU), in a petri dish at 28.5^0^C incubator. Adult *Tg(fli1a:EGFP)* zebrafish were maintained in a 14 hour light/10 hour dark cycle in a recirculating water system at 28^0^C. OMM2.5 cells were prepared for transplantation as described in previously published report [98]. OMM2.5 cells were labelled with 6 mg/ml Dil (Sigma-Aldrich) stain solution prepared in 1 X PBS for 30 min at 37^0^C. OMM2.5 Dil labelled cells were washed twice with 1 X PBS and resuspended in complete media. OMM2.5 Dil cells were filtered through a 40 µm cell strainer prior to microinjection. Approximately 200 - 500 labelled cells were micro- injected (microINJECTOR^TM^, Tritech Research; Los Angeles, CA, USA) into the perivitelline space of two days old *Tg(fli1a:EGFP)* zebrafish larvae under anesthesia (0.05 mg/ml MS222; Sigma-Aldrich). Larvae were imaged using a fluorescent microscope (SMZ1500 attached with DS-Fi2 camera head, Nikon; Tokyo, Japan) for red fluorescence and placed individually into 48-well plates. Only larvae with tumor cells correctly implanted in the perivitelline space were included in the study. Larvae (0 days post treatment (dpt)) were treated with either 0.5% DMSO, 20 µM ACY-1215 (MCE, MedChemExpress; NJ, USA), 20 µM Dacarbazine (TCI, Tokyo Chemical Industry; Tokyo, Japan) or 1.25 µM ML329 for three days at 35^0^C and imaged at both perivitelline space and caudal vein plexus post treatment (25 - 32 pooled larvae were used per treatment group). Differences to transplanted OMM2.5 Dil labelled cells primary fluorescence between 0 dpt and 3 dpt was measured, normalized and calculated using ImageJ. Before drug treatment, toxicity assays were performed with either 0.5% DMSO, ACY-1215, Dacarbazine or ML329 (ranges from 1 - 100 µM). Eight larvae (4 larvae/well) per treatment group was exposed to the desired concentration of drug solutions for 3 days in 24/48-well plates at 35^0^C and imaged at 3 dpt. One-way ANOVA with Dunnett’s Test for Multiple Comparisons or Student’s T test statistical analysis was performed using GraphPad Prism.

### Proteome profiling and mass spectrometry analysis

OMM2.5 cells were seeded at a density of 1 x 10^6^ cells per well and drug treated for 4 or 24 hours with 0.5% DMSO or 20 µM ACY-1215, in duplicate (N = 4). Protein was isolated using PreOmics iST For protein/proteomics preparation kit (PreOmics GmbH; Martinsried, Germany) according to manufacturer’s protocol. Mass Spectrometry and bioinformatic analysis of samples were performed as described previously [99]. Slight variations to methodology consist of raw data processing performed with MaxQuant v1.6.10.43, with MS/MS spectra and database search performed against Uniprot *Homo sapiens* database (2020_05) containing 75,074 entries [99]. Pathway analysis of enriched proteins (a fold change of (+/-) ≥ 1.2 and a *p* value of ≤ 0.05) was performed using ClueGo (v2.5.8) [100] and Cluepedia (v1.5.8) [101] plugins in Cytoscape (v3.8.2) [102] with the *Homo sapiens* (9606) marker set. GO: Biological Process functional pathway databases, consisting of 18058 genes, were used. GO tree levels (min = 3; max = 8) and GO term restriction (min#genes = 3, min% = 4%) were set and terms were grouped using a Kappa Score Threshold of 0.4. The classification was performed by two-sided hypergeometric enrichment test, and its probability value was corrected by the Benjamini-Hochberg method (Adjusted % Term p-value < 0.05).

### Western blot analysis

Protein was isolated from Mel270, Mel285, OMM2.5 and ARPE19 cells at a cell density of 1 x 10^6^ or 4 x 10^5^ and immunoblotting was performed as described [38]. For validation of proteomics data, protein isolated for MS study was utilized. Protein concentrations were measured using BCA protein assay kit (ThermoFisher Scientific; Waltham, MA, United States) in accordance with manufacturer’s instructions and 10 μg of protein was loaded per lane (N = 3 - 4). Blots were probed for HDAC6 (1:1000; #7558, Cell Signaling Technology; Danvers, MA, USA, kindly provided by Dr. Tríona Ní Chonghaile, Dublin, Ireland), acetylated α-tubulin (1:1000; #5335, Cell Signaling Technology), MITF (1:1000; #ab122982, Abcam kindly provided by Dr. Desmond Tobin, Dublin, Ireland), cleaved PARP (1:1000; #5625, Cell Signaling Technology kindly provided by Dr. Emma Dorris, Dublin, Ireland), p-ERK (1:500; #sc-7383, Santa Cruz Biotechnology Inc.; Dallas, TX, USA), ERK (1:500; #sc-514302, Santa Cruz Biotechnology, Inc.), α-tubulin (1:1000; #sc- 5286, Santa Cruz Biotechnology Inc.), GAPDH (1:1000; #2118, Cell Signaling Technology) and β-actin (1:1000; #A5441, Sigma-Aldrich). Anti-rabbit IgG, HRP-linked Antibody (1:3000; #7074s, Cell Signaling Technology) and anti-mouse IgG, HRP-linked Antibody (1:3000; #7076s, Cell Signaling Technology) were used as secondary antibodies. Signal was detected with enhanced chemiluminescence substrate (Pierce™ ECL Western Blotting Substrate; ThermoFisher Scientific). Densitometry analysis was performed using ImageJ and One-way ANOVA with Dunnett’s Test for Multiple Comparisons or Student’s T test statistical analysis using GraphPad Prism.

### Flow cytometry analysis

A total of 300,000 OMM2.5 cells were seeded and treated with 0.5% DMSO, 50 µM Etoposide (Sigma-Aldrich; kindly provided by Dr. William Watson, Dublin, Ireland), 10, 20 and 50 µM ACY-1215 or 20 µM Dacarbazine, in duplicate (N = 3 - 4) for 4, 24 and 96 hours at 37^0^C with 5% CO_2_. Cells were trypsinized and filtered through 50 µm CellTrics filter. Live cells were labelled sequentially with YO-PRO™-1 Iodide (Molecular Probes^TM^ by ThermoFisher Scientific) for 15 mins and Propidium Iodide (PI, Molecular Probes^TM^ by ThermoFisher Scientific) for 3 mins, in the dark at RT to analyze apoptotic events. For cell cycle analysis, cells were fixed in ice cold 70% ethanol at 4^0^C. Cells post-fixed were labelled with 1.25 µl of 1 mg/ml propidium iodide stock and co-treated with 2.5 µl of 10 mg/ml RNase A enzyme (ThermoFisher Scientific) for 30 mins at RT, in the dark. All samples were run on a BD Accuri^TM^ C6 Flow Cytometer (BD Biosciences; NJ, USA) and up to 50,000 events were recorded per sample (N = 3 - 4). YO-PRO™-1 Iodide and PI were excited using a 488 nm laser, and its fluorescence was collected using FL-1 channel (B530/30 band pass filter) and FL-3 channel (B675LP band pass filter), respectively. For cell cycle analysis, PI was excited using a 488 nm laser and its fluorescence collected using FL-2 channel (575/25 band pass filter). Collected samples were gated based on controls (DMSO/Etoposide) and preliminarily analyzed using CFlow Plus Software (v1.0.264.21; BD Biosciences). Reanalysis was performed using FCS Express^TM^ De Novo (Research Edition) v6 software. Instrument was calibrated with manufacturer’s specifications prior to use. Two-Way ANOVA followed by Tukey’s multiple comparison test or Dunnett’s Test for Multiple Comparisons statistical analysis were performed using GraphPad Prism. Detailed information on flow cytometry experiments and analysis performed are provided in **Table S3**.

### TCGA analysis

Survival analyses were performed with package “survminer”, R v3.5.0 (R Foundation for Statistical Computing, Vienna, Austria). Gene expression and clinical data from 80 primary UM included in The Cancer Genome Atlas (TCGA) were collected from the cBioPortal. RNA-seq data were downloaded in Fragments Per Kilobase of exon per million fragments Mapped (FPKM) and then converted to log2 scale. The associations between gene expression and prognosis were assessed by Cox proportional hazard regression models. Progression Free Survival (PFS) and Overall Survival (OS) were used as end points. For categorization of the gene expression into “High” and “Low” categories, median values were used as cut- off. Survival probabilities were plotted on a Kaplan-Meier curve and a Log-rank test was used to compare the two groups. Progression free survival is defined as time to metastatic recurrence. Overall survival is defined as death by any cause.

## Conclusion

This research provides evidence that HDAC6 inhibitors and MITF inhibitors should be considered and further investigated as a potential treatment option for MUM. Specifically, this study proves the efficacy of ACY-1215 as an anti-cancer agent for MUM *in vitro* and *in vivo*. We have additionally elucidated that ACY-1215 regulates MITF expression via p-ERK signaling when used at high concentrations.

**Supplementary Materials**: **Figure S1:** HDAC6 inhibitors present with anti-cancer activity in UM (Mel285, Mel270) and MUM (OMM2.5) cell lines. **Figure S2:** Toxicity effects of ACY-1215 and Dacarbazine in zebrafish larvae. **Figure S3:** Raw Western blot images for HDAC6 expression in UM and MUM cells. **Figure S4:** Potential off-target effects of ACY-1215 on HDAC isozymes. **Figure S5:** Proteome profile of OMM2.5 cells treated with ACY-1215 for 4 hours. **Figure S6:** Pathway analysis map of down- and up-regulated proteins following ACY-1215 treatment for 24 hours. **Figure S7:** Raw Western blot images for acetylated α-tubulin, MITF, p-ERK, ERK and cleaved PARP expression in ACY-1215 treated OMM2.5 cells. **Figure S8:** DNA ploidy in OMM2.5 cells. **Figure S9:** 4 hours ACY-1215 treatment did not have a profound effect on apoptosis pathway in OMM2.5 cells. **Figure S10:** Toxicity screen of ML329 in zebrafish larvae. **Table S1:** List of downregulated proteins and associated pathways after 24 hours of ACY-1215 treatment. **Table S2:** List of upregulated proteins and associated pathways following ACY-1215 treatment for 24 hours. **Table S3:** Minimum Information about a Flow Cytometry Experiment (MIFlowCyt).

## Acknowledgments

We sincerely thank: Prof. Jacintha O’Sullivan for her time and advice on clonogenic assay methodology and imaging of the plates; Assoc. Prof. Alfonso Blanco Fernández from UCD Conway Flow Cytometry Core and Dr. Eugene Dillon from UCD Conway Mass Spectrometry Core facilities for their technical expertise, advice and training in operation of equipment and data analysis. We thank Patient Advocate Melody Buckley and Ocular Melanoma Ireland for their support and contribution to our lay abstract and UM Awareness campaign activities.

## Author Contributions

Conceptualization, H.S. and B.N.K.; Methodology, H.S., S.GM., S.M., J.M.P., W.W., D.J.T., L.D.J. and B.N.K.; Validation, H.S., S.GM., K.S., J.M.P., W.W., D.J.T., L.D.J. and B.N.K.; Formal Analysis, H.S. and B.N.K.; Investigation, H.S. and S.GM.; Resources, H.S., W.W., D.J.T., L.D.J., and B.N.K; Data Curation, H.S.; Writing - Original Draft Preparation, H.S.; Writing - Review & Editing, H.S., S.G.M., S.M., K.S., J.M.P., W.W., D.J.T., L.D.J. and B.N.K.; Visualization, H.S.; Supervision, B.N.K.; Project Administration, H.S. and B.N.K.; Funding Acquisition, H.S. and B.N.K. All authors have reviewed and approved the submitted version of the manuscript.

## Data Availability Statement

Access to raw datasets will be provided upon request to corresponding author.

## References

1. Grisanti, S.; Tura, A. Uveal Melanoma. In Noncutaneous Melanoma, Scott, J.F., Gerstenblith, M.R., Eds.; Brisbane (AU), 2018.

2. Jager, M.J.; Shields, C.L.; Cebulla, C.M.; Abdel-Rahman, M.H.; Grossniklaus, H.E.; Stern, M.H.; Carvajal, R.D.; Belfort, R.N.; Jia, R.; Shields, J.A.;, et al. Uveal melanoma. Nat Rev Dis Primers 2020, 6, 24, doi:10.1038/s41572-020-0158-0.

3. Baily, C.; O’Neill, V.; Dunne, M.; Cunningham, M.; Gullo, G.; Kennedy, S.; Walsh, P.M.; Deady, S.; Horgan, N. Uveal Melanoma in Ireland. Ocul Oncol Pathol 2019, 5, 195–204, doi:10.1159/000492391.

4. Krantz, B.A.; Dave, N.; Komatsubara, K.M.; Marr, B.P.; Carvajal, R.D. Uveal melanoma: epidemiology, etiology, and treatment of primary disease. Clin Ophthalmol 2017, 11, 279–289, doi:10.2147/OPTH.S89591.

5. Kaliki, S.; Shields, C.L. Uveal melanoma: relatively rare but deadly cancer. Eye (Lond) 2017, 31, 241–257, doi:10.1038/eye.2016.275.

6. Yang, J.; Manson, D.K.; Marr, B.P.; Carvajal, R.D. Treatment of uveal melanoma: where are we now? Ther Adv Med Oncol 2018, 10, 1758834018757175, doi:10.1177/1758834018757175.

7. Slater, K.; Hoo, P.S.; Buckley, A.M.; Piulats, J.M.; Villanueva, A.; Portela, A.; Kennedy, B.N. Evaluation of oncogenic cysteinyl leukotriene receptor 2 as a therapeutic target for uveal melanoma. Cancer Metastasis Rev 2018, 37, 335–345, doi:10.1007/s10555-018-9751-z.

8. Carvajal, R.D.; Schwartz, G.K.; Tezel, T.; Marr, B.; Francis, J.H.; Nathan, P.D. Metastatic disease from uveal melanoma: treatment options and future prospects. Br J Ophthalmol 2017, 101, 38–44, doi:10.1136/bjophthalmol-2016-309034.

9. Rodriguez-Vidal, C.; Fernandez-Diaz, D.; Fernandez-Marta, B.; Lago-Baameiro, N.; Pardo, M.; Silva, P.; Paniagua, L.; Blanco-Teijeiro, M.J.; Pineiro, A.; Bande, M. Treatment of Metastatic Uveal Melanoma: Systematic Review. Cancers (Basel) 2020, 12, doi:10.3390/cancers12092557.

10. Carvajal, R.D.; Piperno-Neumann, S.; Kapiteijn, E.; Chapman, P.B.; Frank, S.; Joshua, A.M.; Piulats, J.M.; Wolter, P.; Cocquyt, V.; Chmielowski, B.;, et al. Selumetinib in Combination With Dacarbazine in Patients With Metastatic Uveal Melanoma: A Phase III, Multicenter, Randomized Trial (SUMIT). J Clin Oncol 2018, 36, 1232–1239, doi:10.1200/JCO.2017.74.1090.

11. O’Neill, P.A.; Butt, M.; Eswar, C.V.; Gillis, P.; Marshall, E. A prospective single arm phase II study of dacarbazine and treosulfan as first-line therapy in metastatic uveal melanoma. Melanoma Res 2006, 16, 245–248, doi:10.1097/01.cmr.0000205017.38859.07.

12. Schinzari, G.; Rossi, E.; Cassano, A.; Dadduzio, V.; Quirino, M.; Pagliara, M.; Blasi, M.A.; Barone, C. Cisplatin, dacarbazine and vinblastine as first line chemotherapy for liver metastatic uveal melanoma in the era of immunotherapy: a single institution phase II study. Melanoma Res 2017, 27, 591–595, doi:10.1097/CMR.0000000000000401.

13. Woodman, S.E. Metastatic uveal melanoma: biology and emerging treatments. Cancer J 2012, 18, 148–152, doi:10.1097/PPO.0b013e31824bd256.

14. Chua, V.; Mattei, J.; Han, A.; Johnston, L.; LiPira, K.; Selig, S.M.; Carvajal, R.D.; Aplin, A.E.; Patel, S.P. The Latest on Uveal Melanoma Research and Clinical Trials: Updates from the Cure Ocular Melanoma (CURE OM) Science Meeting (2019). Clin Cancer Res 2021, 27, 28–33, doi:10.1158/1078-0432.CCR-20-2536.

15. Pandiani, C.; Beranger, G.E.; Leclerc, J.; Ballotti, R.; Bertolotto, C. Focus on cutaneous and uveal melanoma specificities. Genes Dev 2017, 31, 724–743, doi:10.1101/gad.296962.117.

16. Nathan, P.; Hassel, J.C.; Rutkowski, P.; Baurain, J.F.; Butler, M.O.; Schlaak, M.; Sullivan, R.J.; Ochsenreither, S.; Dummer, R.; Kirkwood, J.M.; et al. Overall Survival Benefit with Tebentafusp in Metastatic Uveal Melanoma. N Engl J Med 2021, 385, 1196–1206, doi:10.1056/NEJMoa2103485.

17. Aldana-Masangkay, G.I.; Sakamoto, K.M. The role of HDAC6 in cancer. J Biomed Biotechnol 2011, 2011, 875824, doi:10.1155/2011/875824.

18. Eckschlager, T.; Plch, J.; Stiborova, M.; Hrabeta, J. Histone Deacetylase Inhibitors as Anticancer Drugs. Int J Mol Sci 2017, 18, doi:10.3390/ijms18071414.

19. McClure, J.J.; Li, X.; Chou, C.J. Advances and Challenges of HDAC Inhibitors in Cancer Therapeutics. Adv Cancer Res 2018, 138, 183–211, doi:10.1016/bs.acr.2018.02.006.

20. West, A.C.; Johnstone, R.W. New and emerging HDAC inhibitors for cancer treatment. J Clin Invest 2014, 124, 30–39, doi:10.1172/JCI69738.

21. Jenke, R.; Ressing, N.; Hansen, F.K.; Aigner, A.; Buch, T. Anticancer Therapy with HDAC Inhibitors: Mechanism-Based Combination Strategies and Future Perspectives. Cancers (Basel) 2021, 13, doi:10.3390/cancers13040634.

22. Lu, X.; Ning, Z.; Li, Z.; Cao, H.; Wang, X. Development of chidamide for peripheral T-cell lymphoma, the first orphan drug approved in China. Intractable Rare Dis Res 2016, 5, 185–191, doi:10.5582/irdr.2016.01024.

23. Moschos, M.M.; Dettoraki, M.; Androudi, S.; Kalogeropoulos, D.; Lavaris, A.; Garmpis, N.; Damaskos, C.; Garmpi, A.; Tsatsos, M. The Role of Histone Deacetylase Inhibitors in Uveal Melanoma: Current Evidence. Anticancer Res 2018, 38, 3817–3824, doi:10.21873/anticanres.12665.

24. Jespersen, H.; Olofsson Bagge, R.; Ullenhag, G.; Carneiro, A.; Helgadottir, H.; Ljuslinder, I.; Levin, M.; All-Eriksson, C.; Andersson, B.; Stierner, U.;, et al. Concomitant use of pembrolizumab and entinostat in adult patients with metastatic uveal melanoma (PEMDAC study): protocol for a multicenter phase II open label study. BMC Cancer 2019, 19, 415, doi:10.1186/s12885-019-5623-3.

25. Ny, L.; Jespersen, H.; Karlsson, J.; Alsen, S.; Filges, S.; All-Eriksson, C.; Andersson, B.; Carneiro, A.; Helgadottir, H.; Levin, M.;, et al. The PEMDAC phase 2 study of pembrolizumab and entinostat in patients with metastatic uveal melanoma. Nat Commun 2021, 12, 5155, doi:10.1038/s41467-021-25332-w.

26. Nencetti, S.; Cuffaro, D.; Nuti, E.; Ciccone, L.; Rossello, A.; Fabbi, M.; Ballante, F.; Ortore, G.; Carbotti, G.; Campelli, F.;, et al. Identification of histone deacetylase inhibitors with (arylidene)aminoxy scaffold active in uveal melanoma cell lines. J Enzyme Inhib Med Chem 2021, 36, 34–47, doi:10.1080/14756366.2020.1835883.

27. Amengual, J.E.; Lue, J.K.; Ma, H.; Lichtenstein, R.; Shah, B.; Cremers, S.; Jones, S.; Sawas, A. First-in-Class Selective HDAC6 Inhibitor (ACY-1215) Has a Highly Favorable Safety Profile in Patients with Relapsed and Refractory Lymphoma. Oncologist 2021, 26, 184–e366, doi:10.1002/onco.13673.

28. Lee, E.K.; Tan-Wasielewski, Z.; Matulonis, U.A.; Birrer, M.J.; Wright, A.A.; Horowitz, N.; Konstantinopoulos, P.A.; Curtis, J.; Liu, J.F. Results of an abbreviated Phase Ib study of the HDAC6 inhibitor ricolinostat and paclitaxel in recurrent ovarian, fallopian tube, or primary peritoneal cancer. Gynecol Oncol Rep 2019, 29, 118–122, doi:10.1016/j.gore.2019.07.010.

29. Vogl, D.T.; Raje, N.; Jagannath, S.; Richardson, P.; Hari, P.; Orlowski, R.; Supko, J.G.; Tamang, D.; Yang, M.; Jones, S.S.;, et al. Ricolinostat, the First Selective Histone Deacetylase 6 Inhibitor, in Combination with Bortezomib and Dexamethasone for Relapsed or Refractory Multiple Myeloma. Clin Cancer Res 2017, 23, 3307–3315, doi:10.1158/1078-0432.CCR-16-2526.

30. Li, T.; Zhang, C.; Hassan, S.; Liu, X.; Song, F.; Chen, K.; Zhang, W.; Yang, J. Histone deacetylase 6 in cancer. J Hematol Oncol 2018, 11, 111, doi:10.1186/s13045-018-0654-9.

31. Boyault, C.; Sadoul, K.; Pabion, M.; Khochbin, S. HDAC6, at the crossroads between cytoskeleton and cell signaling by acetylation and ubiquitination. Oncogene 2007, 26, 5468–5476, doi:10.1038/sj.onc.1210614.

32. Auzmendi-Iriarte, J.; Saenz-Antonanzas, A.; Mikelez-Alonso, I.; Carrasco-Garcia, E.; Tellaetxe-Abete, M.; Lawrie, C.H.; Sampron, N.; Cortajarena, A.L.; Matheu, A. Characterization of a new small-molecule inhibitor of HDAC6 in glioblastoma. Cell Death Dis 2020, 11, 417, doi:10.1038/s41419-020-2586-x.

33. Moufarrij, S.; Srivastava, A.; Gomez, S.; Hadley, M.; Palmer, E.; Austin, P.T.; Chisholm, S.; Roche, K.; Yu, A.; Li, J.;, et al. Combining DNMT and HDAC6 inhibitors increases anti-tumor immune signaling and decreases tumor burden in ovarian cancer. Sci Rep 2020, 10, 3470, doi:10.1038/s41598-020-60409-4.

34. Kuroki, H.; Anraku, T.; Kazama, A.; Shirono, Y.; Bilim, V.; Tomita, Y. Histone deacetylase 6 inhibition in urothelial cancer as a potential new strategy for cancer treatment. Oncol Lett 2021, 21, 64, doi:10.3892/ol.2020.12315.

35. Jager, M.J.; Magner, J.A.; Ksander, B.R.; Dubovy, S.R. Uveal Melanoma Cell Lines: Where do they come from? (An American Ophthalmological Society Thesis). Trans Am Ophthalmol Soc 2016, 114, T5.

36. Chen, P.W.; Murray, T.G.; Uno, T.; Salgaller, M.L.; Reddy, R.; Ksander, B.R. Expression of MAGE genes in ocular melanoma during progression from primary to metastatic disease. Clin Exp Metastasis 1997, 15, 509-518, doi:10.1023/a:1018479011340.

37. Ksander, B.R.; Rubsamen, P.E.; Olsen, K.R.; Cousins, S.W.; Streilein, J.W. Studies of tumorinfiltrating lymphocytes from a human choroidal melanoma. Invest Ophthalmol Vis Sci 1991, 32, 3198-3208.

38. Slater, K.; Heeran, A.B.; Garcia-Mulero, S.; Kalirai, H.; Sanz-Pamplona, R.; Rahman, A.; Al-Attar, N.; Helmi, M.; O’Connell, F.; Bosch, R.;, et al. High Cysteinyl Leukotriene Receptor 1 Expression Correlates with Poor Survival of Uveal Melanoma Patients and Cognate Antagonist Drugs Modulate the Growth, Cancer Secretome, and Metabolism of Uveal Melanoma Cells. Cancers (Basel) 2020, 12, doi:10.3390/cancers12102950.

39. Depetter, Y.; Geurs, S.; De Vreese, R.; Goethals, S.; Vandoorn, E.; Laevens, A.; Steenbrugge, J.; Meyer, E.; de Tullio, P.; Bracke, M.;, et al. Selective pharmacological inhibitors of HDAC6 reveal biochemical activity but functional tolerance in cancer models. Int J Cancer 2019, 145, 735–747, doi:10.1002/ijc.32169.

40. Cosenza, M.; Civallero, M.; Marcheselli, L.; Sacchi, S.; Pozzi, S. Ricolinostat, a selective HDAC6 inhibitor, shows anti-lymphoma cell activity alone and in combination with bendamustine. Apoptosis 2017, 22, 827–840, doi:10.1007/s10495-017-1364-4.

41. Tan, Y.; Zhang, S.; Zhu, H.; Chu, Y.; Zhou, H.; Liu, D.; Huo, J. Histone deacetylase 6 selective inhibitor ACY1215 inhibits cell proliferation and enhances the chemotherapeutic effect of 5-fluorouracil in HCT116 cells. Ann Transl Med 2019, 7, 2, doi:10.21037/atm.2018.11.48.

42. Hartman, M.L.; Czyz, M. Pro-survival role of MITF in melanoma. J Invest Dermatol 2015, 135, 352–358, doi:10.1038/jid.2014.319.

43. Kim, N.; Kim, S.; Lee, M.W.; Jeon, H.J.; Ryu, H.; Kim, J.M.; Lee, H.J. MITF Promotes Cell Growth, Migration and Invasion in Clear Cell Renal Cell Carcinoma by Activating the RhoA/YAP Signal Pathway. Cancers (Basel) 2021, 13, doi:10.3390/cancers13122920.

44. Loercher, A.E.; Tank, E.M.; Delston, R.B.; Harbour, J.W. MITF links differentiation with cell cycle arrest in melanocytes by transcriptional activation of INK4A. J Cell Biol 2005, 168, 35–40, doi:10.1083/jcb.200410115.

45. Mooy, C.; Vissers, K.; Luyten, G.; Mulder, A.; Stijnen, T.; de Jong, P.; Bosman, F. DNA flow cytometry in uveal melanoma: the effect of pre-enucleation irradiation. Br J Ophthalmol 1995, 79, 174–177, doi:10.1136/bjo.79.2.174.

46. Ehlers, J.P.; Worley, L.; Onken, M.D.; Harbour, J.W. Integrative genomic analysis of aneuploidy in uveal melanoma. Clin Cancer Res 2008, 14, 115–122, doi:10.1158/1078-0432.CCR-07-1825.

47. Faloon, P.W.; Bennion, M.; Weiner, W.S.; Smith, R.A.; Wurst, J.; Weiwer, M.; Hartland, C.; Mosher, C.M.; Johnston, S.; Porubsky, P.;, et al. A Small Molecule Inhibitor of the MITF Molecular Pathway. In Probe Reports from the NIH Molecular Libraries Program; Bethesda (MD), 2010.

48. Yee, A.J.; Bensinger, W.I.; Supko, J.G.; Voorhees, P.M.; Berdeja, J.G.; Richardson, P.G.; Libby, E.N.; Wallace, E.E.; Birrer, N.E.; Burke, J.N.;, et al. Ricolinostat plus lenalidomide, and dexamethasone in relapsed or refractory multiple myeloma: a multicentre phase 1b trial. Lancet Oncol 2016, 17, 1569–1578, doi:10.1016/S1470-2045(16)30375-8.

49. Levinzon, L.; Madigan, M.; Nguyen, V.; Hasic, E.; Conway, M.; Cherepanoff, S. Tumour Expression of Histone Deacetylases in Uveal Melanoma. Ocul Oncol Pathol 2019, 5, 153–161, doi:10.1159/000490038.

50. Putcha, P.; Yu, J.; Rodriguez-Barrueco, R.; Saucedo-Cuevas, L.; Villagrasa, P.; Murga-Penas, E.; Quayle, S.N.; Yang, M.; Castro, V.; Llobet-Navas, D.;, et al. HDAC6 activity is a non-oncogene addiction hub for inflammatory breast cancers. Breast Cancer Res 2015, 17, 149, doi:10.1186/s13058-015-0658-0.

51. Chen, X.; Li, Y.; Yao, T.; Jia, R. Benefits of Zebrafish Xenograft Models in Cancer Research. Front Cell Dev Biol 2021, 9, 616551, doi:10.3389/fcell.2021.616551.

52. Gamble, J.T.; Elson, D.J.; Greenwood, J.A.; Tanguay, R.L.; Kolluri, S.K. The Zebrafish Xenograft Models for Investigating Cancer and Cancer Therapeutics. Biology (Basel) 2021, 10, doi:10.3390/biology10040252.

53. Fazio, M.; Ablain, J.; Chuan, Y.; Langenau, D.M.; Zon, L.I. Zebrafish patient avatars in cancer biology and precision cancer therapy. Nat Rev Cancer 2020, 20, 263–273, doi:10.1038/s41568-020-0252-3.

54. van der Ent, W.; Burrello, C.; Teunisse, A.F.; Ksander, B.R.; van der Velden, P.A.; Jager, M.J.; Jochemsen, A.G.; Snaar-Jagalska, B.E. Modeling of human uveal melanoma in zebrafish xenograft embryos. Invest Ophthalmol Vis Sci 2014, 55, 6612–6622, doi:10.1167/iovs.14-15202.

55. Lin, A.; Giuliano, C.J.; Palladino, A.; John, K.M.; Abramowicz, C.; Yuan, M.L.; Sausville, E.L.; Lukow, D.A.; Liu, L.; Chait, A.R.;, et al. Off-target toxicity is a common mechanism of action of cancer drugs undergoing clinical trials. Sci Transl Med 2019, 11, doi:10.1126/scitranslmed.aaw8412.

56. Ali, A.; Zhang, F.; Maguire, A.; Byrne, T.; Weiner-Gorzel, K.; Bridgett, S.; O’Toole, S.; O’Leary, J.; Beggan, C.; Fitzpatrick, P.; et al. HDAC6 Degradation Inhibits the Growth of High-Grade Serous Ovarian Cancer Cells. Cancers (Basel) 2020, 12, doi:10.3390/cancers12123734.

57. Wloga, D.; Joachimiak, E.; Fabczak, H. Tubulin Post-Translational Modifications and Microtubule Dynamics. Int J Mol Sci 2017, 18, doi:10.3390/ijms18102207.

58. Yokoyama, S.; Feige, E.; Poling, L.L.; Levy, C.; Widlund, H.R.; Khaled, M.; Kung, A.L.; Fisher, D.E. Pharmacologic suppression of MITF expression via HDAC inhibitors in the melanocyte lineage. Pigment Cell Melanoma Res 2008, 21, 457–463, doi:10.1111/j.1755-148X.2008.00480.x.

59. Simmons, J.L.; Pierce, C.J.; Al-Ejeh, F.; Boyle, G.M. MITF and BRN2 contribute to metastatic growth after dissemination of melanoma. Sci Rep 2017, 7, 10909, doi:10.1038/s41598-017-11366-y.

60. Vachtenheim, J. The Many Roles of MITF in Melanoma. Single-Cell Biology 2017, 6, doi:10.4172/2168-9431.1000162.

61. Yajima, I.; Kumasaka, M.Y.; Thang, N.D.; Goto, Y.; Takeda, K.; Iida, M.; Ohgami, N.; Tamura, H.; Yamanoshita, O.; Kawamoto, Y.;, et al. Molecular Network Associated with MITF in Skin Melanoma Development and Progression. J Skin Cancer 2011, 2011, 730170, doi:10.1155/2011/730170.

62. Kawakami, A.; Fisher, D.E. The master role of microphthalmia-associated transcription factor in melanocyte and melanoma biology. Lab Invest 2017, 97, 649–656, doi:10.1038/labinvest.2017.9.

63. Goding, C.R.; Arnheiter, H. MITF-the first 25 years. Genes Dev 2019, 33, 983–1007, doi:10.1101/gad.324657.119.

64. Vachtenheim, J.; Borovansky, J. “Transcription physiology” of pigment formation in melanocytes: central role of MITF. Exp Dermatol 2010, 19, 617–627, doi:10.1111/j.1600-0625.2009.01053.x.

65. D’Mello, S.A.; Finlay, G.J.; Baguley, B.C.; Askarian-Amiri, M.E. Signaling Pathways in Melanogenesis. Int J Mol Sci 2016, 17, doi:10.3390/ijms17071144.

66. Shibahara, S.; Takeda, K.; Yasumoto, K.; Udono, T.; Watanabe, K.; Saito, H.; Takahashi, K. Microphthalmia-associated transcription factor (MITF): multiplicity in structure, function, and regulation. J Investig Dermatol Symp Proc 2001, 6, 99–104, doi:10.1046/j.0022-202x.2001.00010.x.

67. Levy, C.; Khaled, M.; Fisher, D.E. MITF: master regulator of melanocyte development and melanoma oncogene. Trends Mol Med 2006, 12, 406–414, doi:10.1016/j.molmed.2006.07.008.

68. Garraway, L.A.; Widlund, H.R.; Rubin, M.A.; Getz, G.; Berger, A.J.; Ramaswamy, S.; Beroukhim, R.; Milner, D.A.; Granter, S.R.; Du, J.;, et al. Integrative genomic analyses identify MITF as a lineage survival oncogene amplified in malignant melanoma. Nature 2005, 436, 117–122, doi:10.1038/nature03664.

69. Vachtenheim, J.; Ondrusova, L. Microphthalmia-associated transcription factor expression levels in melanoma cells contribute to cell invasion and proliferation. Exp Dermatol 2015, 24, 481–484, doi:10.1111/exd.12724.

70. Aida, S.; Sonobe, Y.; Yuhki, M.; Sakata, K.; Fujii, T.; Sakamoto, H.; Mizuno, T. MITF suppression by CH5552074 inhibits cell growth in melanoma cells. Cancer Chemother Pharmacol 2017, 79, 1187–1193, doi:10.1007/s00280-017-3317-6.

71. Wiedemann, G.M.; Aithal, C.; Kraechan, A.; Heise, C.; Cadilha, B.L.; Zhang, J.; Duewell, P.; Ballotti, R.; Endres, S.; Bertolotto, C.;, et al. Microphthalmia-Associated Transcription Factor (MITF) Regulates Immune Cell Migration into Melanoma. Transl Oncol 2019, 12, 350–360, doi:10.1016/j.tranon.2018.10.014.

72. Hsiao, Y.J.; Chang, W.H.; Chen, H.Y.; Hsu, Y.C.; Chiu, S.C.; Chiang, C.C.; Chang, G.C.; Chen, Y.J.; Wang, C.Y.; Chen, Y.M.;, et al. MITF functions as a tumor suppressor in non-small cell lung cancer beyond the canonically oncogenic role. Aging (Albany NY) 2020, 13, 646–674, doi:10.18632/aging.202171.

73. Guhan, S.M.; Artomov, M.; McCormick, S.; Njauw, C.; Stratigos, A.J.; Shannon, K.; Ellisen, L.W.; Tsao, H. Cancer risks associated with the germline MITF(E318K) variant. Sci Rep 2020, 10, 17051, doi:10.1038/s41598-020-74237-z.

74. Nooron, N.; Ohba, K.; Takeda, K.; Shibahara, S.; Chiabchalard, A. Dysregulated Expression of MITF in Subsets of Hepatocellular Carcinoma and Cholangiocarcinoma. Tohoku J Exp Med 2017, 242, 291–302, doi:10.1620/tjem.242.291.

75. Chen, X.; Wang, J.; Shen, H.; Lu, J.; Li, C.; Hu, D.N.; Dong, X.D.; Yan, D.; Tu, L. Epigenetics, microRNAs, and carcinogenesis: functional role of microRNA-137 in uveal melanoma. Invest Ophthalmol Vis Sci 2011, 52, 1193–1199, doi:10.1167/iovs.10-5272.

76. Lee, D.H.; Won, H.R.; Ryu, H.W.; Han, J.M.; Kwon, S.H. The HDAC6 inhibitor ACY1215 enhances the anticancer activity of oxaliplatin in colorectal cancer cells. Int J Oncol 2018, 53, 844–854, doi:10.3892/ijo.2018.4405.

77. Ryu, H.W.; Shin, D.H.; Lee, D.H.; Won, H.R.; Kwon, S.H. A potent hydroxamic acid-based, small-molecule inhibitor A452 preferentially inhibits HDAC6 activity and induces cytotoxicity toward cancer cells irrespective of p53 status. Carcinogenesis 2018, 39, 72–83, doi:10.1093/carcin/bgx121.

78. Cao, J.; Lv, W.; Wang, L.; Xu, J.; Yuan, P.; Huang, S.; He, Z.; Hu, J. Ricolinostat (ACY-1215) suppresses proliferation and promotes apoptosis in esophageal squamous cell carcinoma via miR- 30d/PI3K/AKT/mTOR and ERK pathways. Cell Death Dis 2018, 9, 817, doi:10.1038/s41419-018-0788-2.

79. Ruan, Y.; Wang, L.; Lu, Y. HDAC6 inhibitor, ACY1215 suppress the proliferation and induce apoptosis of gallbladder cancer cells and increased the chemotherapy effect of gemcitabine and oxaliplatin. Drug Dev Res 2021, 82, 598–604, doi:10.1002/ddr.21780.

80. Boulares, A.H.; Yakovlev, A.G.; Ivanova, V.; Stoica, B.A.; Wang, G.; Iyer, S.; Smulson, M. Role of poly(ADP-ribose) polymerase (PARP) cleavage in apoptosis. Caspase 3-resistant PARP mutant increases rates of apoptosis in transfected cells. J Biol Chem 1999, 274, 22932–22940, doi:10.1074/jbc.274.33.22932.

81. Kaufmann, S.H.; Desnoyers, S.; Ottaviano, Y.; Davidson, N.E.; Poirier, G.G. Specific proteolytic cleavage of poly(ADP-ribose) polymerase: an early marker of chemotherapy-induced apoptosis. Cancer Res 1993, 53, 3976–3985.

82. Wu, J.Y.; Moses, N.; Bai, W.; Zhang, X.M. Implications of the HDAC6-ERK1 feed-forward loop in immunotherapy. J Immunol Sci 2018, 2, 59–68, doi:10.29245/2578-3009/2018/3.1143.

83. Wu, J.Y.; Xiang, S.; Zhang, M.; Fang, B.; Huang, H.; Kwon, O.K.; Zhao, Y.; Yang, Z.; Bai, W.; Bepler, G.;, et al. Histone deacetylase 6 (HDAC6) deacetylates extracellular signal-regulated kinase 1 (ERK1) and thereby stimulates ERK1 activity. J Biol Chem 2018, 293, 1976–1993, doi:10.1074/jbc.M117.795955.

84. Williams, K.A.; Zhang, M.; Xiang, S.; Hu, C.; Wu, J.Y.; Zhang, S.; Ryan, M.; Cox, A.D.; Der, C.J.; Fang, B.;, et al. Extracellular signal-regulated kinase (ERK) phosphorylates histone deacetylase 6 (HDAC6) at serine 1035 to stimulate cell migration. J Biol Chem 2013, 288, 33156–33170, doi:10.1074/jbc.M113.472506.

85. Zhang, S.Z., H.; Zhou, B.; Chu, Y.; Huo, J.; Tan, Y.; and Liu, D.;. Histone deacetylase 6 is overexpressed and promotes tumor growth of colon cancer through regulation of the MAPK/ERK signal pathway. OncoTargets and Therapy 2019, 12, 2409–2419, doi:10.2147/OTT.S194986.

86. Peng, U.; Wang, Z.; Pei, S.; Ou, Y.; Hu, P.; Liu, W.; Song, J. ACY-1215 accelerates vemurafenib induced cell death of BRAF-mutant melanoma cells via induction of ER stress and inhibition of ERK activation. Oncol Rep 2017, *37*, 1270–1276, doi:10.3892/or.2016.5340.

87. Chuang, M.J.; Wu, S.T.; Tang, S.H.; Lai, X.M.; Lai, H.C.; Hsu, K.H.; Sun, K.H.; Sun, G.H.; Chang, S.Y.; Yu, D.S.;, et al. The HDAC inhibitor LBH589 induces ERK-dependent prometaphase arrest in prostate cancer via HDAC6 inactivation and down-regulation. PLoS One 2013, 8, e73401, doi:10.1371/journal.pone.0073401.

88. Lim, J.A.; Juhnn, Y.S. Isoproterenol increases histone deacetylase 6 expression and cell migration by inhibiting ERK signaling via PKA and Epac pathways in human lung cancer cells. Exp Mol Med 2016, 48, e204, doi:10.1038/emm.2015.98.

89. Boru, G.; Cebulla, C.M.; Sample, K.M.; Massengill, J.B.; Davidorf, F.H.; Abdel-Rahman, M.H. Heterogeneity in Mitogen-Activated Protein Kinase (MAPK) Pathway Activation in Uveal Melanoma With Somatic GNAQ and GNA11 Mutations. Invest Ophthalmol Vis Sci 2019, 60, 2474–2480, doi:10.1167/iovs.18-26452.

90. Chen, X.; Wu, Q.; Tan, L.; Porter, D.; Jager, M.J.; Emery, C.; Bastian, B.C. Combined PKC and MEK inhibition in uveal melanoma with GNAQ and GNA11 mutations. Oncogene 2014, 33, 4724–4734, doi:10.1038/onc.2013.418.

91. Steeb, T.; Wessely, A.; Ruzicka, T.; Heppt, M.V.; Berking, C. How to MEK the best of uveal melanoma: A systematic review on the efficacy and safety of MEK inhibitors in metastatic or unresectable uveal melanoma. Eur J Cancer 2018, 103, 41–51, doi:10.1016/j.ejca.2018.08.005.

92. Sagoo, M.S.; Harbour, J.W.; Stebbing, J.; Bowcock, A.M. Combined PKC and MEK inhibition for treating metastatic uveal melanoma. Oncogene 2014, 33, 4722–4723, doi:10.1038/onc.2013.555.

93. Zuidervaart, W.; van Nieuwpoort, F.; Stark, M.; Dijkman, R.; Packer, L.; Borgstein, A.M.; Pavey, S.; van der Velden, P.; Out, C.; Jager, M.J.;, et al. Activation of the MAPK pathway is a common event in uveal melanomas although it rarely occurs through mutation of BRAF or RAS. Br J Cancer 2005, 92, 2032–2038, doi:10.1038/sj.bjc.6602598.

94. Mouriaux, F.; Vincent, S.; Kherrouche, Z.; Maurage, C.A.; Planque, N.; Monte, D.; Labalette, P.; Saule, S. Microphthalmia transcription factor analysis in posterior uveal melanomas. Exp Eye Res 2003, 76, 653–661, doi:10.1016/s0014-4835(03)00082-4.

95. Iwamoto, S.; Burrows, R.C.; Grossniklaus, H.E.; Orcutt, J.; Kalina, R.E.; Boehm, M.; Bothwell, M.A.; Schmidt, R. Immunophenotype of conjunctival melanomas: comparisons with uveal and cutaneous melanomas. Arch Ophthalmol 2002, 120, 1625–1629, doi:10.1001/archopht.120.12.1625.

96. Perrino, C.M.; Wang, J.F.; Collins, B.T. Microphthalmia transcription factor immunohistochemistry for FNA biopsy of ocular malignant melanoma. Cancer Cytopathol 2015, 123, 394–400, doi:10.1002/cncy.21531.

97. Maurya, D.K. ColonyCountJ: A User-Friendly Image J Add-on Program for Quantification of Different Colony Parameters in Clonogenic Assay. Journal of Clinical Toxicology 2017, 7, doi:10.4172/2161-0495.1000358.

98. Rouhi, P.; Jensen, L.D.; Cao, Z.; Hosaka, K.; Lanne, T.; Wahlberg, E.; Steffensen, J.F.; Cao, Y. Hypoxia-induced metastasis model in embryonic zebrafish. Nat Protoc 2010, 5, 1911–1918, doi:10.1038/nprot.2010.150.

99. Sundaramurthi, H.; Roche, S.L.; Grice, G.L.; Moran, A.; Dillion, E.T.; Campiani, G.; Nathan, J.A.; Kennedy, B.N. Selective Histone Deacetylase 6 Inhibitors Restore Cone Photoreceptor Vision or Outer Segment Morphology in Zebrafish and Mouse Models of Retinal Blindness. Front Cell Dev Biol 2020, 8, 689, doi:10.3389/fcell.2020.00689.

100. Bindea, G.; Mlecnik, B.; Hackl, H.; Charoentong, P.; Tosolini, M.; Kirilovsky, A.; Fridman, W.H.; Pages, F.; Trajanoski, Z.; Galon, J. ClueGO: a Cytoscape plug-in to decipher functionally grouped gene ontology and pathway annotation networks. Bioinformatics 2009, 25, 1091–1093, doi:10.1093/bioinformatics/btp101.

101. Bindea, G.; Galon, J.; Mlecnik, B. CluePedia Cytoscape plugin: pathway insights using integrated experimental and in silico data. Bioinformatics 2013, 29, 661–663, doi:10.1093/bioinformatics/btt019.

102. Shannon, P.; Markiel, A.; Ozier, O.; Baliga, N.S.; Wang, J.T.; Ramage, D.; Amin, N.; Schwikowski, B.; Ideker, T. Cytoscape: a software environment for integrated models of biomolecular interaction networks. Genome Res 2003, 13, 2498–2504, doi:10.1101/gr.1239303.

