## Supplementary data for "Inhibition of Metastatic Uveal Melanoma Cell Proliferation by the Histone Deacetylase 6 Inhibitor, Ricolinostat, is Linked to Altered Microphthalmia-associated Transcription Factor and Phospho-ERK Expression"

### Supplementary Figures and Tables

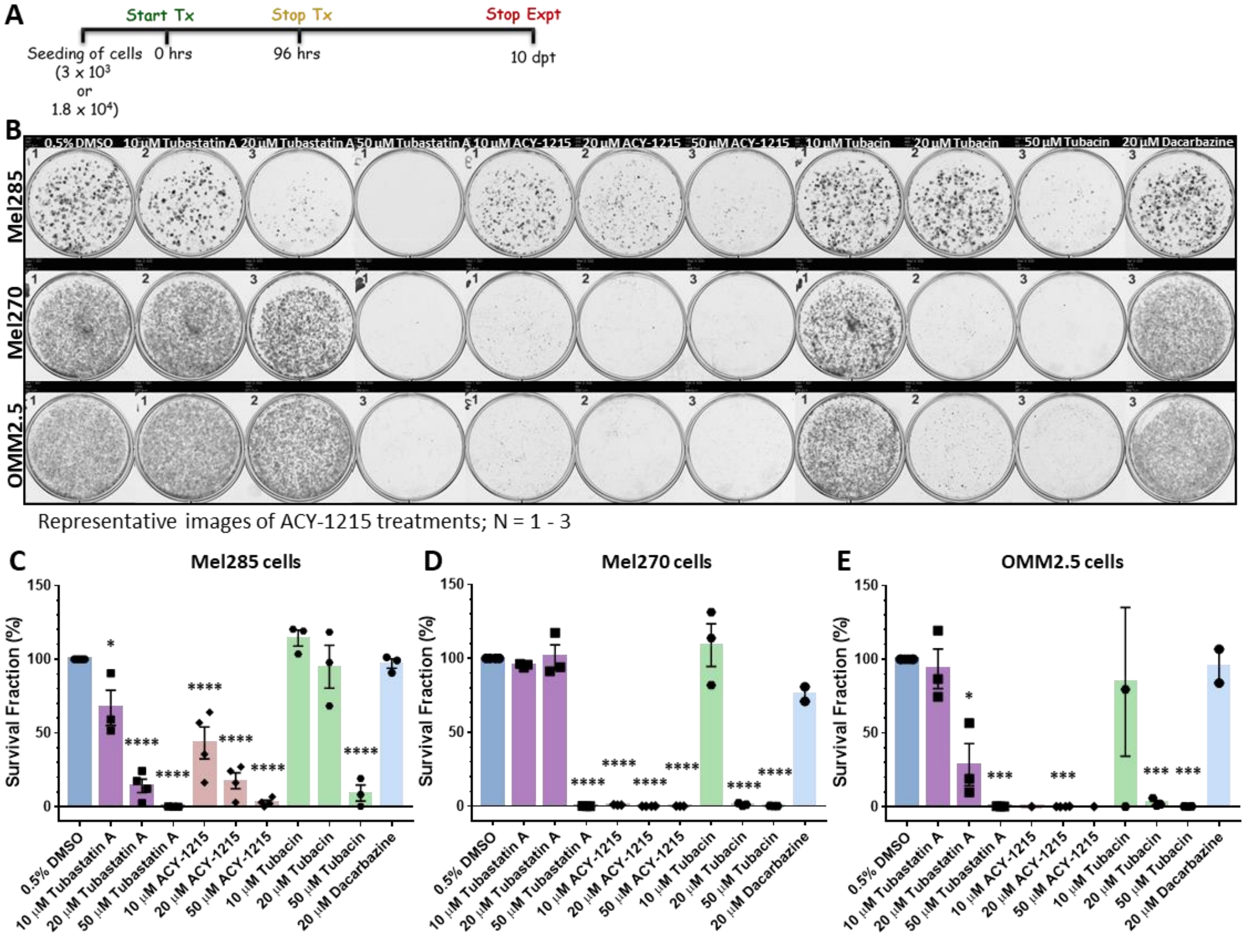


**Figure S1:** HDAC6 inhibitors present with anti-cancer activity in UM (Mel285, Mel270) and MUM (OMM2.5) cell lines. A: Schematic diagram illustrating treatment regime used. B: Representative image of clonogenic assay plates for Mel285 (top panel), Mel270 (middle panel) and OMM2.5 cells (bottom panel) treated with 0.5% DMSO, 10, 20 or 50 µM Tubastatin A, 10, 20 or 50 µM ACY-1215, 10, 20 or 50 µM Tubacin or 20 µM Dacarbazine for 96 hours. C, D and E: A dose-dependent decrease in surviving fraction of colonies was observed across the three different HDAC6i tested in comparison to 0.5% DMSO treatment. Percentage of surviving colonies was comparable between 20 µM Dacarbazine and 0.5% DMSO treatment group. One-way ANOVA with Dunnett's Test for Multiple Comparisons statistical analysis was performed, error bars represent mean + SEM, ****p = 0.0001 (N = 2 - 4).

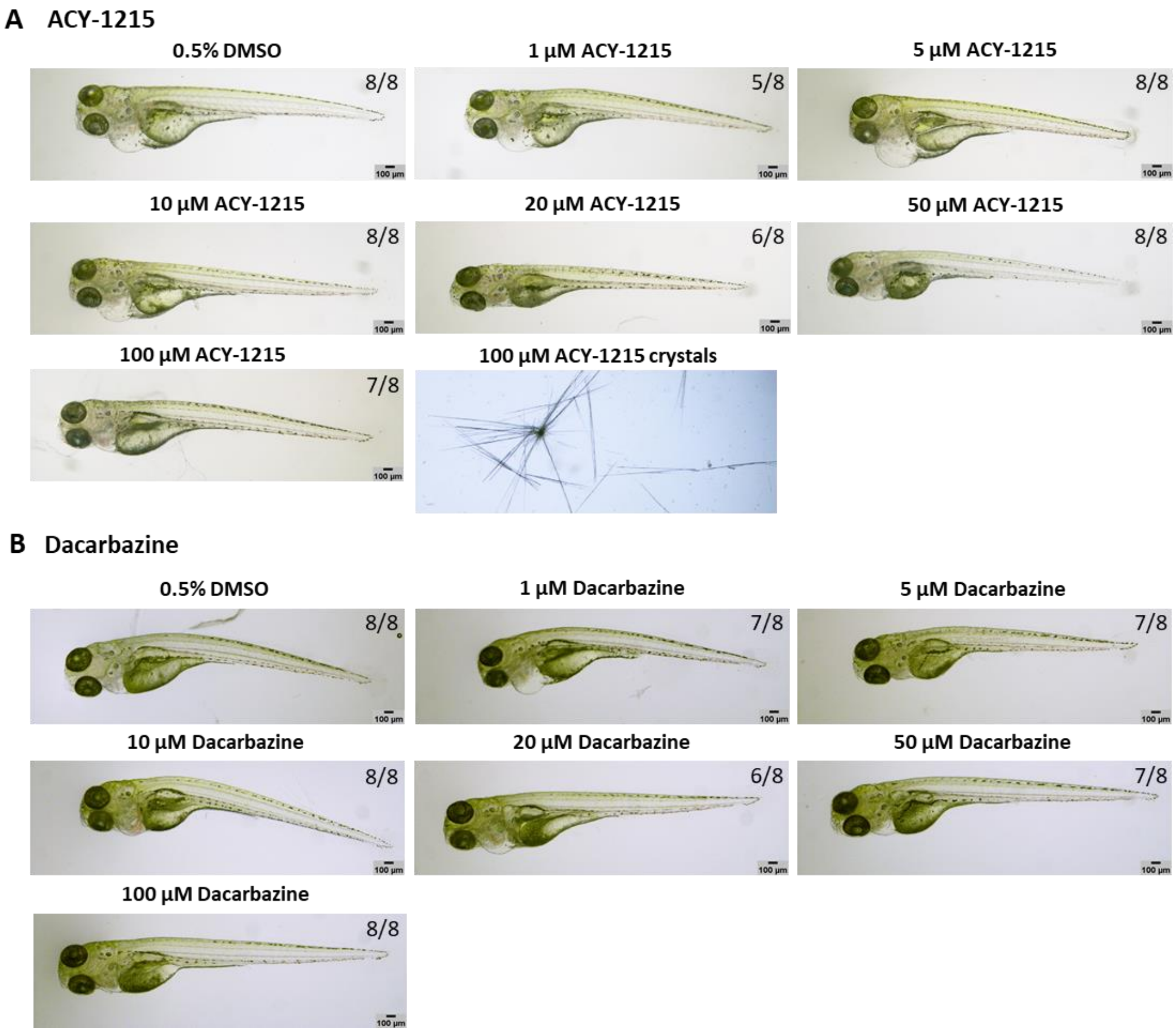


**Figure S2:** Toxicity effects of ACY-1215 and Dacarbazine in zebrafish larvae. A and B: Increasing doses of ACY-1215 and Dacarbazine, up to 100 µM, are well tolerated by 5 days (3 days post treated) old *Tg(fli1a:EGFP)* zebrafish larvae (n = 8 per treatment group). Drug crystals were observed to form at 100 µM ACY-1215.

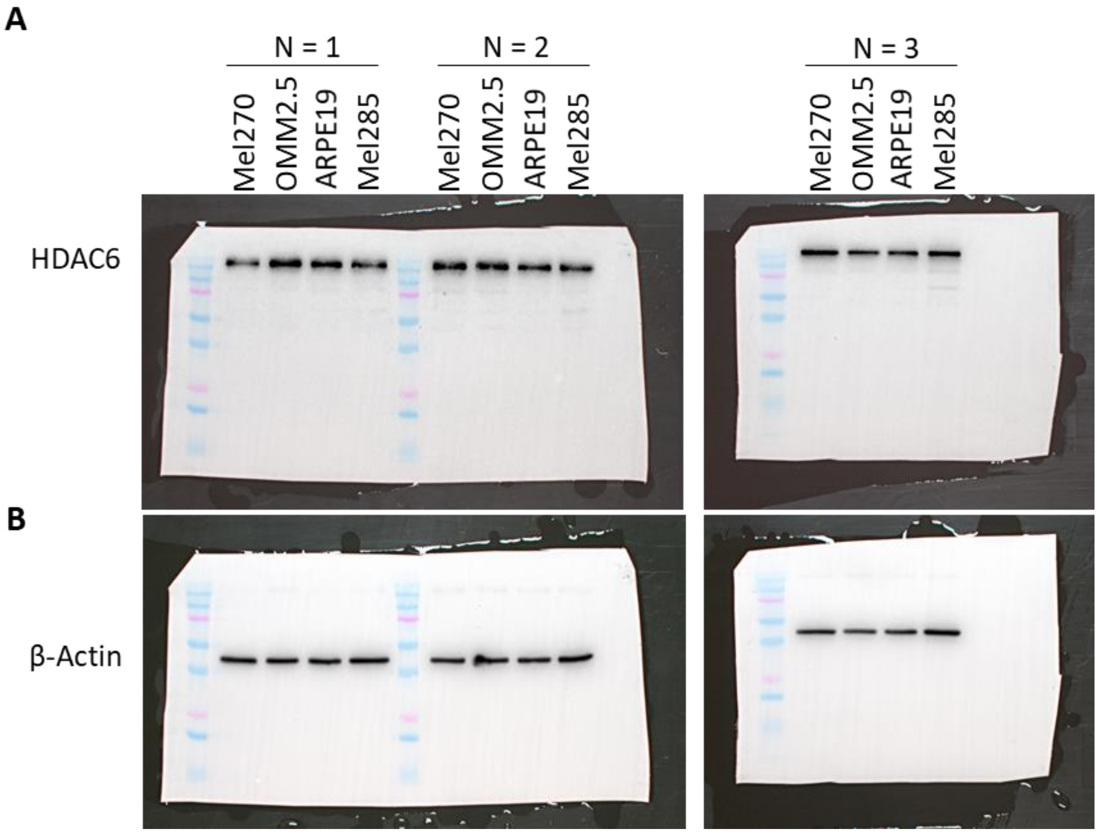


**Figure S3:** Raw Western blot images for HDAC6 expression in UM and MUM cells. A: HDAC6 expression in UM (Mel270, Mel285) and mUM (OMM2.5) cells, is comparable to ARPE19 cells (N = 3). B: β-actin was used as loading control.

**A**

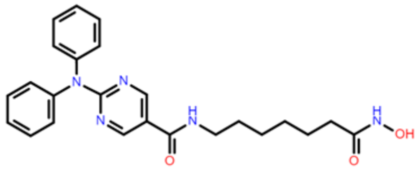

ACY-1215 (Ricolinostat)

**B**

| HDAC | $IC_{50}$ | Activity | HDAC | $IC_{50}$ | Activity |
|---|---|---|---|---|---|
| HDAC6 | 4.7 nM | Potent & Selective | HDAC5 | 5 µM | Minimal |
| HDAC2 | 48 nM | Less Active | HDAC4 | 7 µM | Minimal |
| HDAC3 | 51 nM | Less Active | HDAC9 | > 1 µM | Minimal |
| HDAC1 | 58 nM | Less Active | HDAC11 | > 1 µM | Minimal |
| HDAC8 | 100 nM | Slight | Sirt 1 | > 1 µM | Minimal |
| HDAC7 | 1.4 µM | Minimal | Sirt 2 | > 1 µM | Minimal |

*Adapted from SelleckChem*

**C**

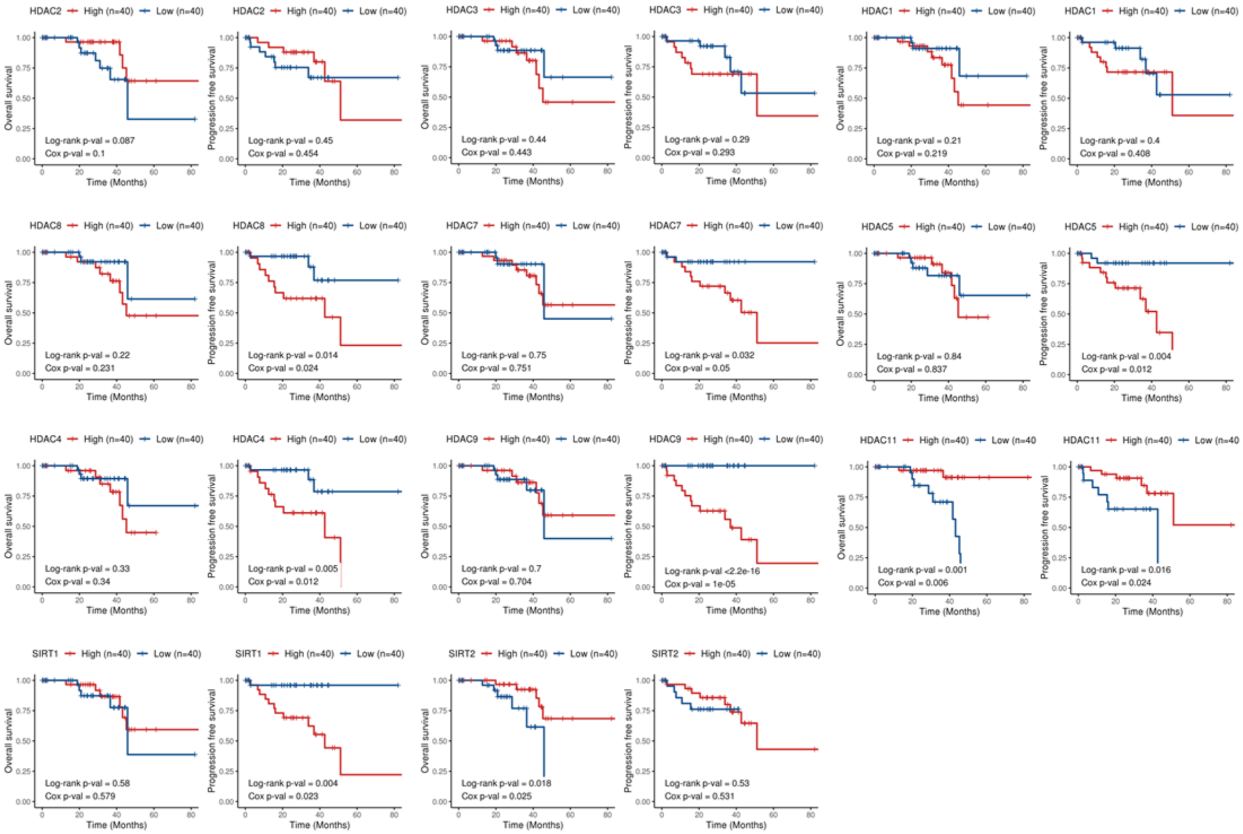


**Figure S4:** Potential off-target effects of ACY-1215 on HDAC isozymes. A: Chemical structure of ACY-1215 drawn using ChemSpider. B: Table highlighting ACY-1215 $IC_{50}$ values against various HDAC isozymes. C: Kaplan-Meier survival curves demonstrating correlation between expression of various HDAC isoforms and overall survival (OS) or progression free survival (PFS) in UM patients. Median values were used as cut-off for high (red) and low (blue) expression levels, with Log-rank *p*-values (categorical variable) and Cox *p*-values (continuous variable) calculated (n = 80).

A

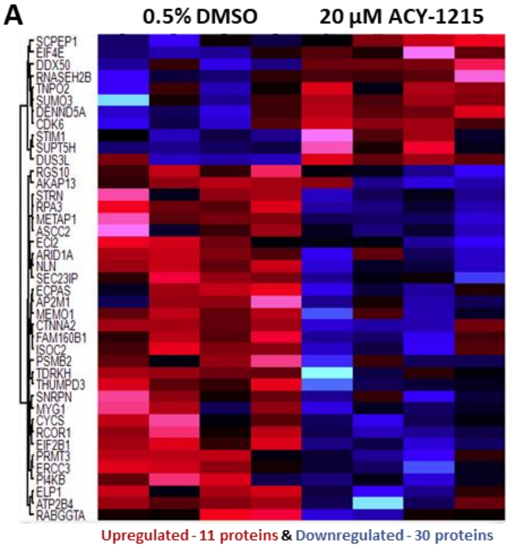


B

| Gene | UNIPROT | Protein Names | -Log Student's T-test p-value | Log Student's T-test Difference | Fold Change |
|---|---|---|---|---|---|
| **Top 10 Downregulated Proteins** | | | | | |
| *CTNNA2* | P26232 | Catenin alpha-2 | 1.78 | - 3.80 | - 13.96 |
| *AKAP13* | Q12802 | A-kinase anchor protein 13 | 1.44 | - 1.90 | - 3.74 |
| *ARID1A* | O14497 | AT-rich interactive domain-containing protein 1A | 1.71 | - 1.86 | - 3.63 |
| *NLN* | Q9BYT8 | Neurolysin, mitochondrial | 2.87 | - 1.54 | - 2.92 |
| *RCOR1* | Q9UKL0 | REST corepressor 1 | 2.31 | - 1.47 | - 2.77 |
| *ELP1* | F5H2T0 | IkappaB kinase complex-associated protein | 1.65 | - 1.37 | - 2.59 |
| *SNRPN* | J3KRY3 | Small nuclear ribonucleoprotein-associated protein N (Fragment) | 1.46 | - 0.82 | - 1.76 |
| *PRMT3* | O60678 | Protein arginine N-methyltransferase 3 | 1.63 | - 0.77 | - 1.71 |
| *ATP2B4* | P23634 | Plasma membrane calcium-transporting ATPase 4 | 1.39 | - 0.69 | - 1.62 |
| *TDRKH* | Q5SZR4 | Tudor and KH domain containing, isoform CRA_a | 1.37 | - 0.62 | - 1.54 |
| **Top 10 Upregulated Proteins** | | | | | |
| **Gene** | **UNIPROT** | **Protein Names** | **-Log Student's T-test p-value** | **Log Student's T-test Difference** | **Fold Change** |
| *DDX50* | A0A087WVC1 | RNA helicase | 2.02 | 1.40 | 2.64 |
| *DUS3L* | Q96G46 | tRNA-dihydrouridine(47) synthase [NAD(P)(+)]-like | 1.62 | 1.16 | 2.24 |
| *SUMO3* | P55854 | Small ubiquitin-related modifier 3 | 1.40 | 1.14 | 2.20 |
| *DENND5A* | H0YDE6 | DENN domain-containing protein 5A (Fragment) | 2.03 | 0.44 | 1.36 |
| *SCPEP1* | Q9HB40 | Retinoid-inducible serine carboxypeptidase | 1.76 | 0.36 | 1.28 |
| *TNPO2* | A0A075B780 | Transportin-2 | 1.35 | 0.32 | 1.25 |
| *SUPT5H* | O00267 | Transcription elongation factor SPT5 | 1.58 | 0.29 | 1.22 |
| *RNASEH2B* | A0A2R8YCX2 | Ribonuclease H2 subunit B | 1.41 | 0.22 | 1.17 |
| *EIF4E* | P06730 | Eukaryotic translation initiation factor 4E | 1.51 | 0.12 | 1.09 |
| *CDK6* | Q00534 | Cyclin-dependent kinase 6 | 1.74 | 0.12 | 1.09 |

C Downregulated proteins - Biological Processes

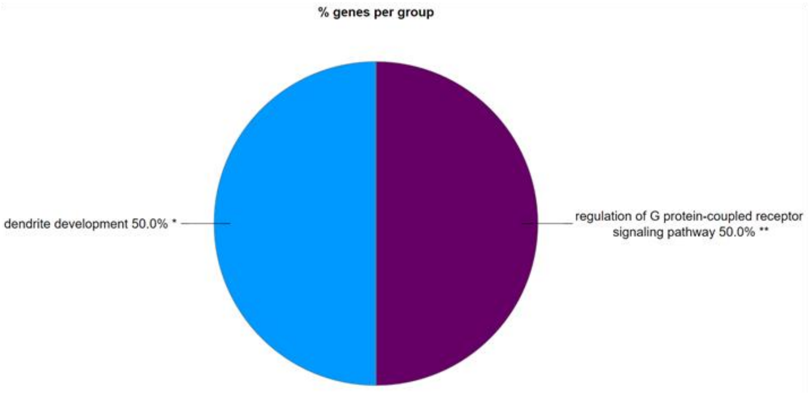


**Figure S5:** Proteome profile of OMM2.5 cells treated with ACY-1215 for 4 hours. A: Heat map presenting significantly differentially expressed proteins after 4 hours of 20 µM ACY-1215 treatment in OMM2.5 cells. 30 (blue) proteins were downregulated, and 11 (red) proteins were upregulated (N = 4). B: Table showing the top 10 down- and up-regulated proteins at 4 hours. C: Enriched protein pathway analysis for GO term: biological processes, identified dendrite development and regulation of G protein-coupled receptor signaling pathway to be significantly downregulated.

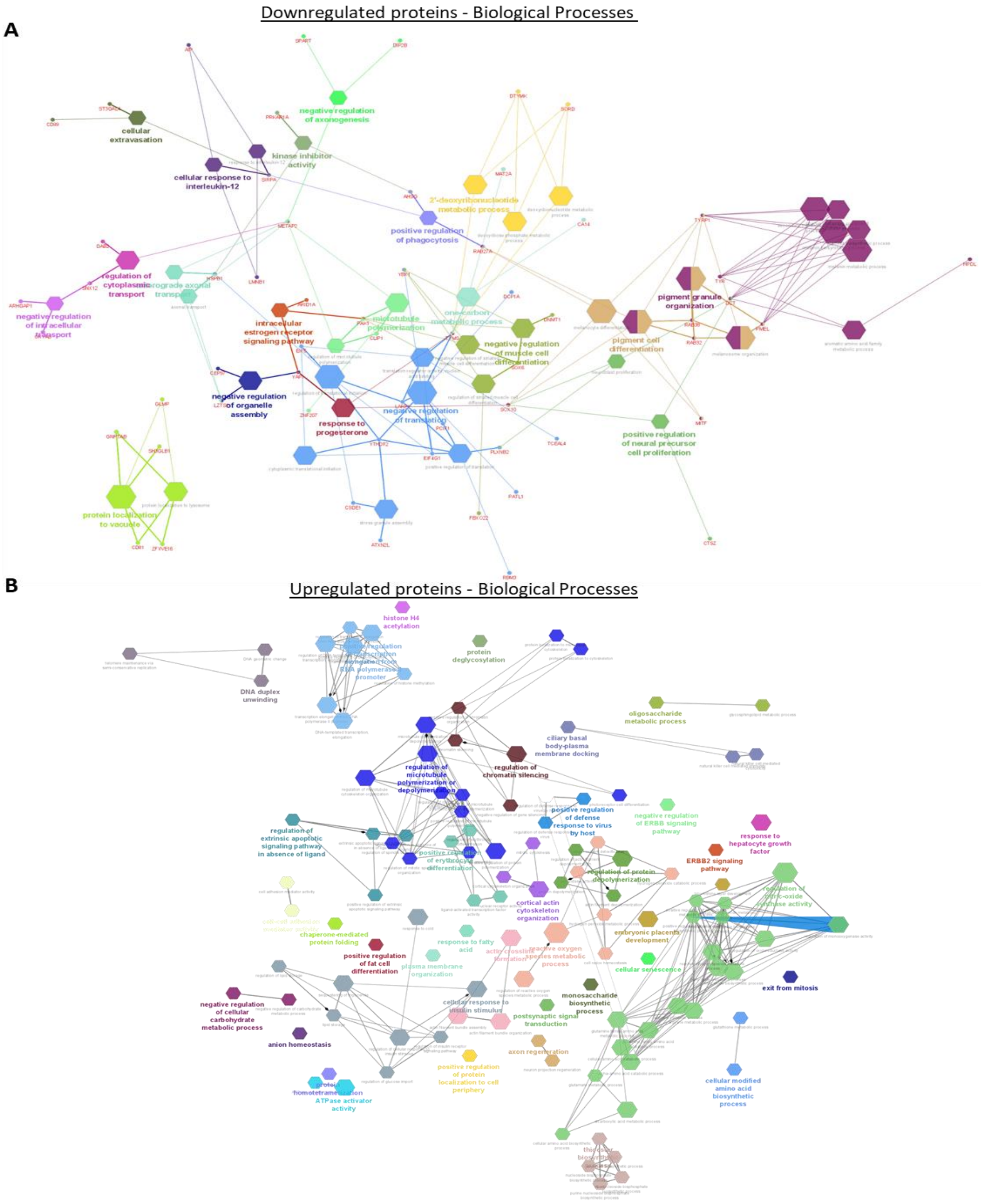


**Figure S6:** GO: Biological processes pathway analysis map of down- and up-regulated proteins following ACY-1215 treatment for 24 hours.

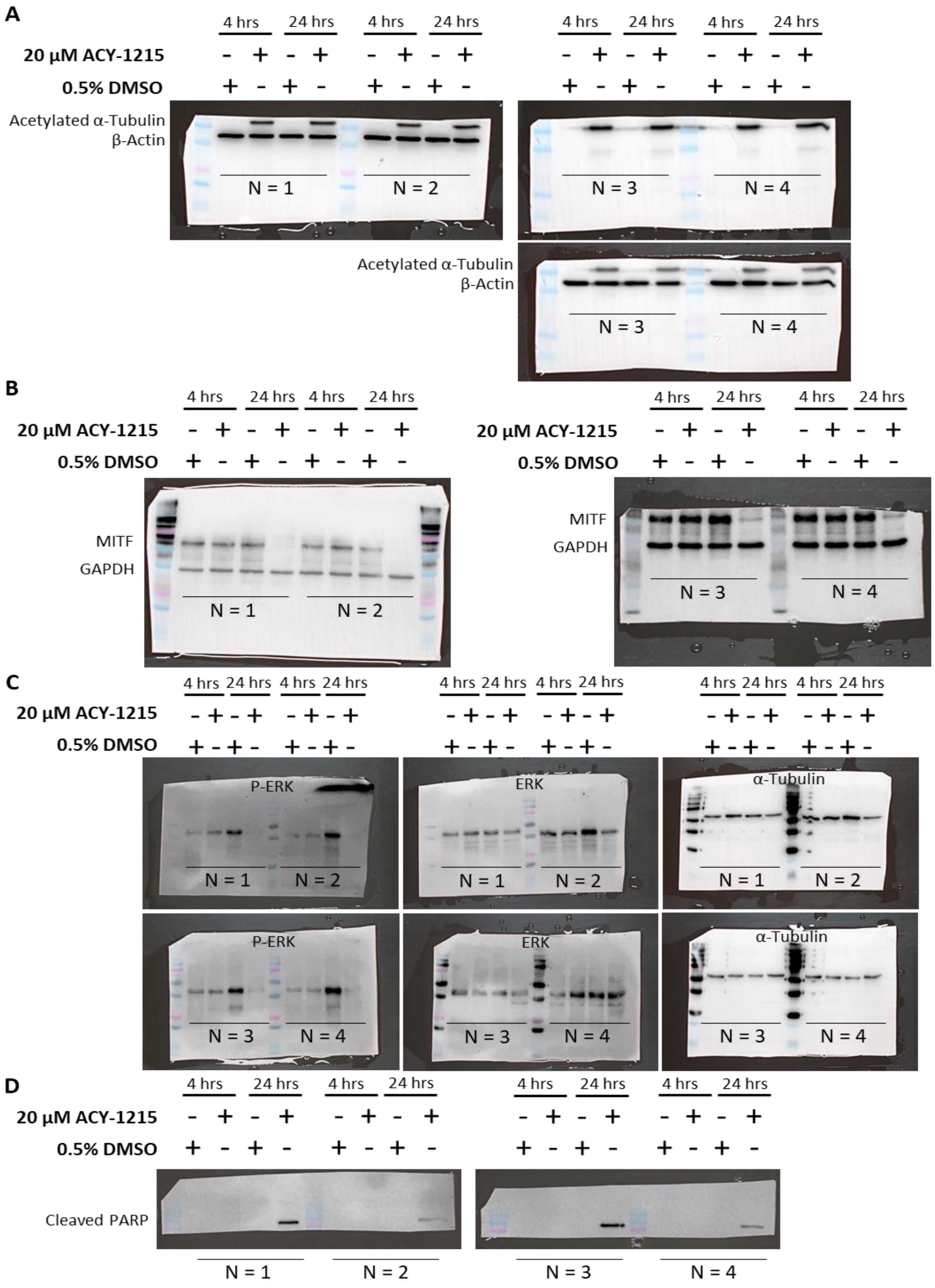
A
4 hrs
24 hrs
20 μM ACY-1215
0.5% DMSO
Acetylated α-Tubulin
β-Actin
N = 1
N = 2
N = 3
N = 4
B
MITF
GAPDH
C
P-ERK
ERK
α-Tubulin
D
Cleaved PARP

**Figure S7:** Raw Western blot images for acetylated α-tubulin, MITF, p-ERK, ERK and cleaved PARP expression in ACY-1215 treated OMM2.5 cells. A: Acetylated α-tubulin expression is upregulated after 4 and 24 hours of 20 μM ACY-1215 treatment in OMM2.5 cells. B: MITF expression remained unchanged after 4 hours of 20 μM ACY-1215 treatment. 24 hours of 20 μM ACY-1215 resulted in a significant reduction in MITF expression levels compared to vehicle control. C: p-ERK/Total ERK expression levels not altered following 4 hours of treatment with 20 μM ACY-1215. 20 μM ACY-1215 treatment for 24 hours led to a significant reduction in p-ERK expression levels, in comparison to vehicle control. D: Cleaved PARP expression was upregulated in OMM2.5 cells 24 hours post treated with 20 μM ACY-1215. β-actin, α-tubulin and GAPDH were used as loading control (N = 4).

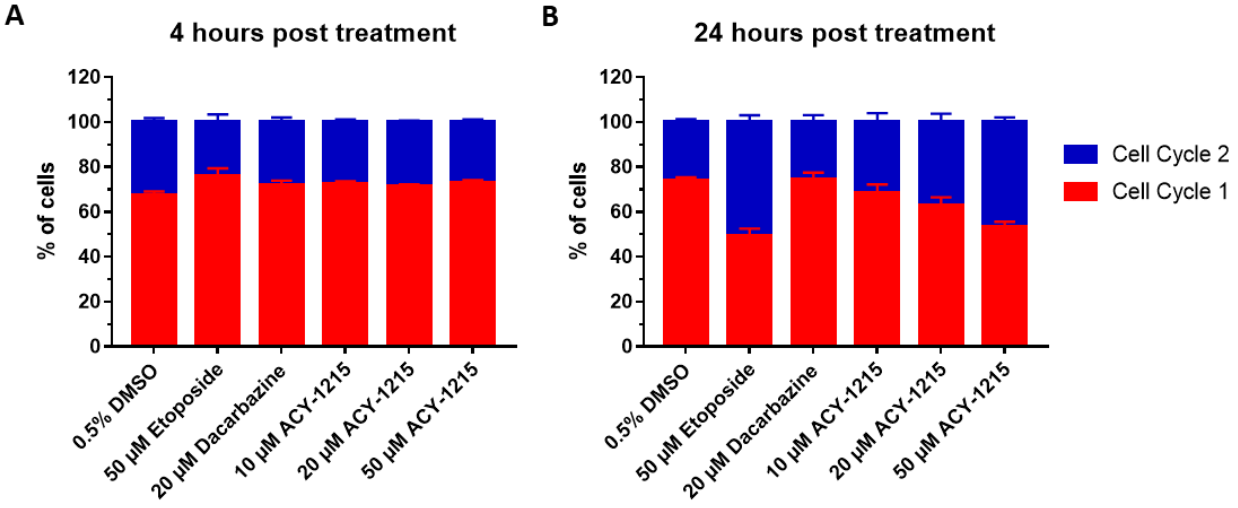


**Figure S8:** DNA ploidy in OMM2.5 cells. A and B: OMM2.5 cells display DNA ploidy as presented by the two cell cycle populations observed (N = 4). Error bars indicate mean <u>+</u> SEM (N = 3)

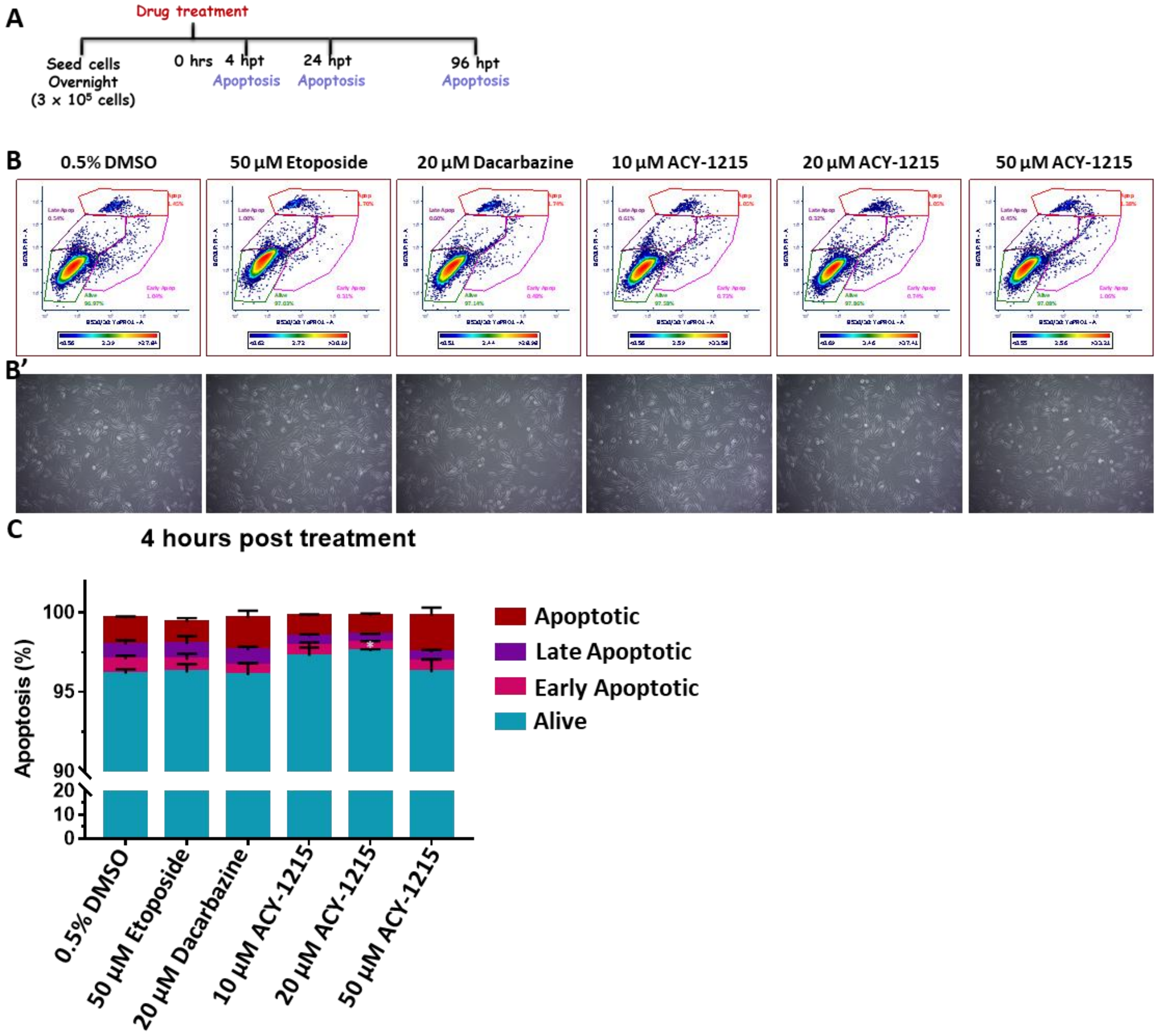


**Figure S9:** 4 hours ACY-1215 treatment did not have a profound effect on apoptosis pathway in OMM2.5 cells. A: Illustration of treatment regime. B: Representative plots of treated OMM2.5 cell singlets gating in different stages of apoptosis. B': Micrographs corresponding to OMM2.5 drug treatment groups at 4 hours. C: No change was observed in live, early, late and apoptotic cell populations following treatment with Etoposide and all concentrations of ACY-1215 treatment. On average, a significant increase in percentage of live cells was detected following 20 µM ACY-1215 (*, $p = 0.03$) treatment compared to 0.5% DMSO treatment. Two-way ANOVA followed by Tukey's Multiple Comparisons statistical test was performed, error bars show mean $\pm$ SEM (N = 3).

### Supplementary Fig 10: Toxicity screen in zebrafish

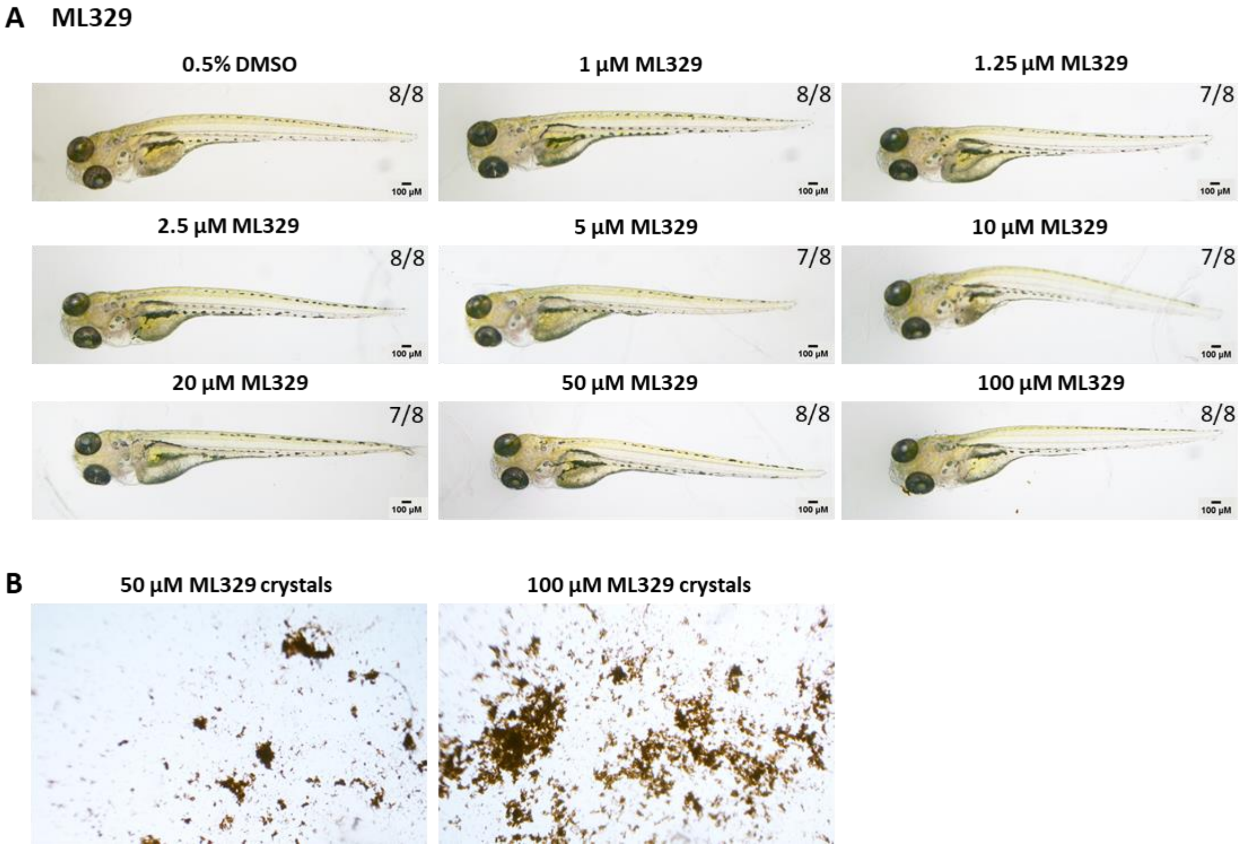


**Figure S10:** Toxicity screen of ML329 in zebrafish larvae. A: Increasing doses of ML329, up to 100 µM, was well tolerated by 5 days (3 days post treated) old *Tg(fli1a:EGFP)* zebrafish larvae (n = 8 per treatment group). B: Drug precipitates observed in embryo media at 50 and 100 µM ML329 concentrations 3 days post treatment.

**Table S1:** List of downregulated proteins and associated pathways after 24 hours of ACY-1215 treatment.

| ID | Term | Associated Genes Found |
|---|---|---|
| GO:0019210 | Kinase inhibitor activity | AHSG, HSPB1, PRKAR1A |
| GO:0050771 | Negative regulation of axonogenesis | DIP2B, METAP2, SPART |
| GO:0006730 | One-carbon metabolic process | CA14, MAT2A, TYMS |
| GO:1902116 | Negative regulation of organelle assembly | CEP97, LZTS1, YAP1 |
| GO:1903649 | Regulation of cytoplasmic transport | DAB2, METAP2, SNX12 |
| GO:0030520 | Intracellular estrogen receptor signaling pathway | ARID1A, PAK1, YAP1 |
| GO:0032387 | Negative regulation of intracellular transport | ARHGAP1, CRYAB, SNX12 |
| GO:0032570 | Response to progesterone | SOX10, TYMS, YAP1 |
| GO:0045123 | Cellular extravasation | CD99, SIRPA, ST3GAL4 |
| GO:0050766 | Positive regulation of phagocytosis | AHSG, RAB27A, SIRPA |
| GO:0070671 | Response to interleukin-12 | AIP, LMNB1, SIRPA |
| GO:0071349 | Cellular response to interleukin-12 | AIP, LMNB1, SIRPA |
| GO:0072665 | Protein localization to vacuole | CD81, GLMP, GNPTAB, SH3GLB1, ZFYVE16 |
| GO:0061462 | Protein localization to lysosome | CD81, GLMP, GNPTAB, ZFYVE16 |
| GO:0098930 | Axonal transport | HSPB1, LZTS1, METAP2 |
| GO:0008089 | Anterograde axonal transport | HSPB1, LZTS1, METAP2 |
| GO:2000179 | Positive regulation of neural precursor cell proliferation | CTSZ, DCT, SOX10 |
| GO:0007405 | Neuroblast proliferation | DCT, PLXNB2, SOX10 |
| GO:0046785 | Microtubule polymerization | CLIP1, METAP2, PAK1, ZNF207 |
| GO:0031113 | Regulation of microtubule polymerization | CLIP1, METAP2, PAK1 |
| GO:0019692 | Deoxyribose phosphate metabolic process | DTYMK, SORD, TYMS |
| GO:0009262 | Deoxyribonucleotide metabolic process | DTYMK, SORD, TYMS |
| GO:0009394 | 2'-deoxyribonucleotide metabolic process | DTYMK, SORD, TYMS |
| GO:0051148 | Negative regulation of muscle cell differentiation | DNMT1, PAK1, SOX6, YBX1 |
| GO:0051153 | Regulation of striated muscle cell differentiation | FBXO22, PAK1, SOX6, YBX1 |
| GO:0051154 | Negative regulation of striated muscle cell differentiation | PAK1, SOX6, YBX1 |
| GO:0050931 | Pigment cell differentiation | MITF, RAB27A, RAB32, RAB38, SOX10, TYRP1 |
| GO:0030318 | Melanocyte differentiation | MITF, RAB27A, SOX10, TYRP1 |

| ID | Term | Associated Genes Found |
|---|---|---|
| GO:0048753 | Pigment granule organization | PMEL, RAB32, RAB38, TYRP1 |
| GO:0032438 | Melanosome organization | PMEL, RAB32, RAB38, TYRP1 |
| GO:0034063 | Stress granule assembly | ATXN2L, CSDE1, YTHDF2 |
| GO:0006446 | Regulation of translational initiation | CSDE1, EIF4G1, EIF5, HSPB1, LARP1, THDF2 |
| GO:0002183 | Cytoplasmic translational initiation | EIF4G1, EIF5, YTHDF2 |
| GO:0017148 | Negative regulation of translation | DCP1A, EIF4G1, LARP1, PATL1, PCIF1, TYMS, YBX1, YTHDF2 |
| GO:0045727 | Positive regulation of translation | EIF4G1, LARP1, PCIF1, PLXNB2, RBM3, YTHDF2 |
| GO:0090079 | Translation regulator activity, nucleic acid binding | EIF4G1, EIF5, LARP1, TCEAL4, TYMS |
| GO:0044550 | Secondary metabolite biosynthetic process | DCT, PMEL, TYR, TYRP1 |
| GO:0046148 | Pigment biosynthetic process | DCT, PMEL, TYR, TYRP1 |
| GO:0050931 | Pigment cell differentiation | MITF, RAB27A, RAB32, RAB38, SOX10, TYRP1 |
| GO:0006582 | Melanin metabolic process | DCT, PMEL, TYR, TYRP1 |
| GO:0009072 | Aromatic amino acid family metabolic process | DCT, HPDL, TYR |
| GO:0048753 | Pigment granule organization | PMEL, RAB32, RAB38, TYRP1 |
| GO:0042438 | Melanin biosynthetic process | DCT, PMEL, TYR, TYRP1 |
| GO:0046189 | Phenol-containing compound biosynthetic process | DCT, PMEL, TYR, TYRP1 |
| GO:0032438 | Melanosome organization | PMEL, RAB32, RAB38, TYRP1 |

**Table S2:** List of upregulated proteins and associated pathways following ACY-1215 treatment for 24 hours.

| ID | Term | Associated Genes Found |
|---|---|---|
| Go:0006517 | Protein deglycosylation | MAN2B1, OS9, SEL1L |
| Go:0090398 | Cellular senescence | ARG2, MAP2K1, PAWR |
| Go:0007009 | Plasma membrane organization | AHNAK2, BAIAP2L1, NDRG1, PRDX6, RAB3A, SLC9A3R1 |
| Go:0010458 | Exit from mitosis | ANLN, EPS8, UBE2S |
| Go:0035728 | Response to hepatocyte growth factor | CRK, GCLC, LGMN |
| Go:0038128 | Erbb2 signaling pathway | CDC37, GRB2, HSP90AA1 |
| Go:0043967 | Histone h4 acetylation | HAT1, IWS1, RUVBL1 |
| Go:0045600 | Positive regulation of fat cell differentiation | AAMDC, LCP1, STK4, SYAP1 |
| Go:0046364 | Monosaccharide biosynthetic process | ALDOC, G6PD, GOT1, SDHAF3, STK4 |
| Go:0051289 | Protein homotetramerization | ACOT13, GLS, TK1 |
| Go:0055081 | Anion homeostasis | ENPP1, GLS, SLC9A3R1 |
| Go:0061077 | Chaperone-mediated protein folding | DNAJB1, HSPA1A, SGTB |
| Go:0070542 | Response to fatty acid | CAT, CCN2, LCP1 |
| Go:0098926 | Postsynaptic signal transduction | KPNA2, RGS10, STAT3 |
| Go:1901185 | Negative regulation of erbb signaling pathway | GRB2, LGMN, MVP |
| Go:1904377 | Positive regulation of protein localization to cell periphery | ACSL3, EPB41L2, SORBS1 |
| Go:0009311 | Oligosaccharide metabolic process | GLA, MAN2B1, NEU1 |
| Go:0006687 | Glycosphingolipid metabolic process | ARSA, GLA, NEU1 |
| Go:0031102 | Neuron projection regeneration | CTNNA1, DHFR, MAP2K1 |
| Go:0031103 | Axon regeneration | CTNNA1, DHFR, MAP2K1 |
| Go:0006749 | Glutathione metabolic process | G6PD, GCLC, GGCT |
| Go:0042398 | Cellular modified amino acid biosynthetic process | DHFR, GCLC, GGCT |
| Go:0045912 | Negative regulation of carbohydrate metabolic process | ENPP1, STAT3, STK4 |
| Go:0010677 | Negative regulation of cellular carbohydrate metabolic process | ENPP1, STAT3, STK4 |
| Go:0060590 | Atpase regulator activity | ATP1B1, DNAJB1, RAB3A |
| Go:0001671 | Atpase activator activity | ATP1B1, DNAJB1, RAB3A |
| Go:0001892 | Embryonic placenta development | CCN1, GRB2, MAP2K1, MIA3, STK4 |
| Go:0060711 | Labyrinthine layer development | CCN1, GRB2, MAP2K1 |

| ID | Term | Associated Genes Found |
|---|---|---|
| Go:0098631 | Cell adhesion mediator activity | BAIAP2L1, CXADR, PDLIM5 |
| Go:0098632 | Cell-cell adhesion mediator activity | BAIAP2L1, CXADR, PDLIM5 |
| Go:0000281 | Mitotic cytokinesis | ANLN, ARL3, RACGAP1 |
| Go:0030866 | Cortical actin cytoskeleton organization | ANLN, EPB41L2, LCP1, RACGAP1 |
| Go:0030865 | Cortical cytoskeleton organization | ANLN, EPB41L2, LCP1, RACGAP1 |
| Go:0032201 | Telomere maintenance via semi-conservative replication | BLM, RFC1, RFC4 |
| Go:0032392 | Dna geometric change | BLM, CETN2, MNAT1, RFC4, RUVBL1 |
| Go:0032508 | Dna duplex unwinding | BLM, CETN2, MNAT1, RFC4, RUVBL1 |
| Go:0002228 | Natural killer cell mediated immunity | CRK, PRDX1, TUBB2A, TUBB4B |
| Go:0042267 | Natural killer cell mediated cytotoxicity | CRK, PRDX1, TUBB2A, TUBB4B |
| Go:0097711 | Ciliary basal body-plasma membrane docking | CEP131, CETN2, HSP90AA1, TUBB2A, TUBB4B |
| Go:0050688 | Regulation of defense response to virus | HSP90AA1, SIN3A, TOMM70 |
| Go:0050691 | Regulation of defense response to virus by host | HSP90AA1, SIN3A, TOMM70 |
| Go:0002230 | Positive regulation of defense response to virus by host | HSP90AA1, SIN3A, TOMM70 |
| Go:0051764 | Actin crosslink formation | BAIAP2L1, EPS8, LCP1 |
| Go:0061572 | Actin filament bundle organization | BAIAP2L1, CCN2, EPS8, LCP1, LIMA1, PAWR, SORBS1 |
| Go:0051017 | Actin filament bundle assembly | BAIAP2L1, CCN2, EPS8, LCP1, LIMA1, PAWR, SORBS1 |
| Go:0060969 | Negative regulation of gene silencing | ATAD2, H1-10, STAT3 |
| Go:1905268 | Negative regulation of chromatin organization | ATAD2, H1-10, SIN3A, SUPT6H |
| Go:0006342 | Chromatin silencing | ATAD2, H1-10, HAT1, SIN3A |
| Go:0031935 | Regulation of chromatin silencing | ATAD2, H1-10, SIN3A |
| Go:0038034 | Signal transduction in absence of ligand | CTNNA1, HSPA1A, PPP2R1B |
| Go:0097192 | Extrinsic apoptotic signaling pathway in absence of ligand | CTNNA1, HSPA1A, PPP2R1B |
| Go:2001238 | Positive regulation of extrinsic apoptotic signaling pathway | CTNNA1, PPP2R1B, STK4 |
| Go:2001239 | Regulation of extrinsic apoptotic signaling pathway in absence of ligand | CTNNA1, HSPA1A, PPP2R1B |
| Go:0098531 | Ligand-activated transcription factor activity | MIA3, NR4A1, STAT3 |
| Go:0045646 | Regulation of erythrocyte differentiation | HSPA1A, MIA3, STAT3 |
| Go:0045648 | Positive regulation of erythrocyte differentiation | HSPA1A, MIA3, STAT3 |
| Go:0004879 | Nuclear receptor activity | MIA3, NR4A1, STAT3 |

| ID | Term | Associated Genes Found |
|---|---|---|
| Go:1901879 | Regulation of protein depolymerization | CKAP2, DSTN, EPS8, LIMA1, METAP1 |
| Go:0051261 | Protein depolymerization | CKAP2, DSTN, EPS8, LIMA1, METAP1 |
| Go:0030042 | Actin filament depolymerization | DSTN, EPS8, LIMA1 |
| Go:0030834 | Regulation of actin filament depolymerization | DSTN, EPS8, LIMA1 |
| Go:0035384 | Thioester biosynthetic process | ACSL3, GCDH, PDP1 |
| Go:0033866 | Nucleoside bisphosphate biosynthetic process | ACSL3, GCDH, PDP1 |
| Go:0034033 | Purine nucleoside bisphosphate biosynthetic process | ACSL3, GCDH, PDP1 |
| Go:0034030 | Ribonucleoside bisphosphate biosynthetic process | ACSL3, GCDH, PDP1 |
| Go:0071616 | Acyl-coa biosynthetic process | ACSL3, GCDH, PDP1 |
| Go:0032968 | Positive regulation of transcription elongation from RNA polymerase ii promoter | BRD4, GTF2F1, SUPT6H |
| Go:0006354 | DNA-templated transcription, elongation | BRD4, GTF2F1, HTATSF1, IWS1, MNAT1, SUPT6H |
| Go:0031060 | Regulation of histone methylation | BRD4, IWS1, SUPT6H |
| Go:0032784 | Regulation of DNA-templated transcription, elongation | BRD4, GTF2F1, HTATSF1, SUPT6H |
| Go:0006368 | Transcription elongation from rna polymerase ii promoter | BRD4, GTF2F1, IWS1, MNAT1, SUPT6H |
| Go:0032786 | Positive regulation of dna-templated transcription, elongation | BRD4, GTF2F1, SUPT6H |
| Go:0034243 | Regulation of transcription elongation from rna polymerase ii promoter | BRD4, GTF2F1, SUPT6H |
| Go:0072593 | Reactive oxygen species metabolic process | ARG2, CAT, CBR1, CCN1, CCN2, DHFR, G6PD, GRB2, NQO2, PRDX1, PRDX6, STAT3 |
| Go:0042743 | Hydrogen peroxide metabolic process | CAT, PRDX1, PRDX6, STAT3 |
| Go:0045454 | Cell redox homeostasis | GCLC, PRDX1, PRDX6 |
| Go:0042744 | Hydrogen peroxide catabolic process | CAT, PRDX1, PRDX6 |
| Go:0098869 | Cellular oxidant detoxification | CAT, DHFR, MAPRE2, PRDX1, PRDX6 |
| Go:2000377 | Regulation of reactive oxygen species metabolic process | ARG2, CBR1, DHFR, G6PD, GRB2, NQO2, STAT3 |
| Go:0004601 | Peroxidase activity | CAT, MAPRE2, PRDX1, PRDX6 |
| Go:0009409 | Response to cold | ATP2B1, HSP90AA1, LCP1, VGF |
| Go:0019915 | Lipid storage | EHD1, ENPP1, LCP1, OSBPL8 |
| Go:0010883 | Regulation of lipid storage | EHD1, LCP1, OSBPL8 |
| Go:0030730 | Sequestering of triglyceride | ENPP1, LCP1, OSBPL8 |

| ID | Term | Associated Genes Found |
|---|---|---|
| Go:1900076 | Regulation of cellular response to insulin stimulus | ATP2B1, ENPP1, LCP1, OSBPL8, SORBS1 |
| Go:0032869 | Cellular response to insulin stimulus | ATP2B1, BAIAP2L1, ENPP1, GCLC, GOT1, GRB2, LCP1, OSBPL8, SORBS1, SYAP1 |
| Go:0046626 | Regulation of insulin receptor signaling pathway | BAIAP2L1, ENPP1, OSBPL8, SORBS1 |
| Go:0046324 | Regulation of glucose import | ENPP1, OSBPL8, SORBS1 |
| Go:0031109 | Microtubule polymerization or depolymerization | ARL3, CKAP2, HSPA1A, MAPRE2, METAP1, NDRG1 |
| Go:0070507 | Regulation of microtubule cytoskeleton organization | ARL3, CKAP2, HSPA1A, MAPRE2, METAP1, NDRG1, TACC3 |
| Go:0044380 | Protein localization to cytoskeleton | CEP131, MAPRE2, METAP1 |
| Go:0090224 | Regulation of spindle organization | HSPA1A, NDRG1, TACC3 |
| Go:0031110 | Regulation of microtubule polymerization or depolymerization | ARL3, CKAP2, HSPA1A, MAPRE2, METAP1, NDRG1 |
| Go:0031112 | Positive regulation of microtubule polymerization or depolymerization | ARL3, HSPA1A, NDRG1 |
| Go:0031113 | Regulation of microtubule polymerization | ARL3, HSPA1A, NDRG1 |
| Go:0032273 | Positive regulation of protein polymerization | ARFIP1, ARL3, BAIAP2L1, GRB2, HSP90AA1, HSPA1A, NDRG1 |
| Go:0060236 | Regulation of mitotic spindle organization | HSPA1A, NDRG1, TACC3 |
| Go:0072698 | Protein localization to microtubule cytoskeleton | CEP131, MAPRE2, METAP1 |
| Go:0090307 | Mitotic spindle assembly | HSPA1A, NDRG1, RACGAP1 |
| Go:0046530 | Photoreceptor cell differentiation | ARL3, MAPRE2, STAT3 |
| Go:0031116 | Positive regulation of microtubule polymerization | ARL3, HSPA1A, NDRG1 |
| Go:2001057 | Reactive nitrogen species metabolic process | ARG2, DDAH1, DDAH2, GLA, HSP90AA1 |
| Go:0046209 | Nitric oxide metabolic process | ARG2, DDAH1, DDAH2, GLA, HSP90AA1 |
| Go:0051341 | Regulation of oxidoreductase activity | DDAH1, DDAH2, DHFR, GLA, HSP90AA1, NOSIP, PDP1 |
| Go:0080164 | Regulation of nitric oxide metabolic process | DDAH1, DDAH2, GLA, HSP90AA1 |
| Go:0006809 | Nitric oxide biosynthetic process | ARG2, DDAH1, DDAH2, GLA, HSP90AA1 |
| Go:0032768 | Regulation of monooxygenase activity | DDAH1, DDAH2, DHFR, GLA, HSP90AA1, NOSIP |
| Go:1904407 | Positive regulation of nitric oxide metabolic process | DDAH1, DDAH2, HSP90AA1 |
| Go:0008652 | Cellular amino acid biosynthetic process | DHFR, GLS, GOT1 |
| Go:0009063 | Cellular amino acid catabolic process | ARG2, DDAH1, DDAH2, GCDH, GLS, GOT1, HIBCH |
| Go:0045428 | Regulation of nitric oxide biosynthetic process | DDAH1, DDAH2, GLA, HSP90AA1 |

| ID | Term | Associated Genes Found |
|---|---|---|
| Go:0043648 | Dicarboxylic acid metabolic process | DHFR, GCLC, GLS, GOT1, L2HGDH, SDHA, SDHAF3 |
| Go:0045429 | Positive regulation of nitric oxide biosynthetic process | DDAH1, DDAH2, HSP90AA1 |
| Go:0050999 | Regulation of nitric-oxide synthase activity | DDAH1, DDAH2, DHFR, GLA, HSP90AA1, NOSIP |
| Go:0009064 | Glutamine family amino acid metabolic process | ARG2, DDAH1, DDAH2, GCLC, GLS, GOT1 |
| Go:1901606 | Alpha-amino acid catabolic process | ACADSB, ARG2, DDAH1, DDAH2, GCDH, GLS, GOT1 |
| Go:0006527 | Arginine catabolic process | ARG2, DDAH1, DDAH2 |
| Go:0006525 | Arginine metabolic process | ARG2, DDAH1, DDAH2 |
| Go:0006536 | Glutamate metabolic process | GCLC, GLS, GOT1 |
| Go:0009065 | Glutamine family amino acid catabolic process | ARG2, DDAH1, DDAH2, GLS, GOT1 |

**Table S3:** Minimum Information about a Flow Cytometry Experiment (MIFlowCyt).

| 1. Experiment Overview | Details |
|---|---|
| 1.1. Purpose | To analyze cell cycle and cell death mechanisms of ACY-1215 treated OMM2.5 cells using flow cytometry. |
| 1.2. Keywords | Cell cycle, Apoptosis, Yo-Pro$^{TM}$1, Propidium Iodide |
| 1.3. Experiment Variables | OMM2.5 cells were treated with either 0.5% DMSO, 50 µM Etoposide, 10, 20 and 50 µM ACY-1215 or 20 µM Dacarbazine. Drug treated samples were analyzed at 4, 24 and 96 hours. Experiments were performed in triplicate and quadruplicate. |
| 1.4. Organization Name and Address | Conway Institute, University College Dublin, Belfield, Dublin 4, Ireland |
| 1.5. Primary Contact Name and Email Address | Breandán Kennedy,<br>Husvinee Sundaramurthi, |
| 1.6. Date | October 2020 - December 2020 |
| 1.7. Conclusions | ACY-1215 treatment arrested cell cycle in the S phase by 24 hours post treatment. A time and dose-dependent increase in apoptotic and late-stage apoptotic cells with a decrease in percentage of viable cells was observed in ACY-1215 treated OMM2.5 cells. |
| 1.8. Quality Control Measures | Etoposide was used as a control for apoptosis analysis, Dacarbazine was used as a clinical control to compare against our drug of interest (ACY-1215) and vehicle control was used in each experiment. Experiments were independently performed 3 to 4 times.<br>BD Accuri$^{TM}$ C6 Flow Cytometer was routinely calibrated according to manufacturer's instructions. |
| 1.9. Other Relevant Experiment Information | Not Applicable. |
| **2. Flow Sample/Specimen Details** | |
| **Sample/Specimen Material Description** | **Details** |
| 2.1.1.1. Sample/Specimen Material description | Drug treated live and post-fixed OMM2.5 cells were collected for apoptosis and cell cycle analysis, respectively, as described in Methods section. |
| 2.1.1.2. Biological Sample Source Description<br><br>2.1.1.3. Biological Sample Source Organism Description<br><br>2.1.1.4. Other Relevant Biological Sample Information | Established metastatic UM cell line - OMM2.5 was kindly provided by Dr. Martine Jager, Leiden, The Netherlands (Jager, M.J.; Magner, J.A.; Ksander, B.R.; Dubovy, S.R. Uveal Melanoma Cell Lines: Where do they come from? (An American Ophthalmological Society Thesis). Trans Am Ophthalmol Soc 2016, 114, T5.). |
| 2.1.2. Environmental sample | Not Applicable. |
| 2.1.3. Other Samples | Not Applicable. |
| 2.2 Sample Characteristics | Drug treated OMM2.5 cells were labeled with PI for cell cycle analysis and Yo-Pro1/PI for apoptosis analysis. Cells undergoing |

| | |
|---|---|
| | different stages of apoptosis, alive and dead cells are distinguished based on intensity of fluorescence. |
| 2.3. Sample Treatment Description | Single cell suspensions were prepared after treatment for 4, 24 or 96 hours with desired drug solutions. Samples were divided into 2 aliquots of 300 µl cell suspension solution. One aliquot of live cells was labeled with Yo-Pro1/PI for apoptosis analysis. While the 2$^{nd}$ tube was fixed with 70% ethanol at 4$^{0}$C for a few days. Post-fixed cells were then labelled with PI prior to analysis of cell cycle. A maximum of 50,000 events were recorded per sample for analysis. |
| 2.4. Fluorescence Reagent Description | YO-PRO™-1 Iodide : Molecular Probes™ by Life Technologies<br>Propidium iodide (PI) : Molecular Probes™ by Life Technologies |
| **3. Instrument Details** | |
| 3.1. Instrument Manufacturer | BD Biosciences (https://www.bdbiosciences.com/en-gb) |
| 3.2. Instrument model | BD Accuri™ C6 Flow Cytometer |
| 3.3. Instrument configuration and settings<br><br>3.3.1. Flow Cell and Fluidics<br><br>3.3.2. Light Sources<br><br>3.3.3. Excitation Optics Configuration<br><br>3.3.4. Optical Filters<br><br>3.3.5. Optical Detectors<br><br>3.3.6. Optical Paths | |
| 3.4. Other Relevant Instrument Details | |
| **4. Data Analysis Details** | |
| 4.1. List-mode Data Files | FCS data files can be obtained upon request to corresponding author, post publication. |
| 4.2. Compensation Description | No compensation was applied |
| 4.3. Data Transformation Details | No transformation was applied |
| 4.4. Gating Details<br><br>4.4.1. Gate description<br>4.4.2. Gate statistics<br>4.4.3. Gate boundaries<br>4.4.4. Other Relevant Gate Information | A representative figure and statistical analysis have been provided in Figures X, X and Supplementary Figures X, X. For additional info please refer to Results and Methods text in manuscript. |
